# Temporal Gene Expression in Apical Culms Shows Early Changes in Cell Wall Biosynthesis Genes in Sugarcane

**DOI:** 10.1101/2021.07.01.450585

**Authors:** Guilherme Kenichi Hosaka, Fernando Henrique Correr, Carla Cristina Silva, Danilo Augusto Sforça, Fernanda Zatti Barreto, Thiago Willian Almeida Balsalobre, Asher Pasha, Anete Pereira de Souza, Nicholas James Provart, Monalisa Sampaio Carneiro, Gabriel Rodrigues Alves Margarido

## Abstract

Multiple genes in sugarcane control sucrose accumulation and the biosynthesis of cell wall components; however, it is unclear how these genes are expressed in its apical culms. To better understand this process, we sequenced mRNA from +1 stem internodes collected from four genotypes with different concentrations of soluble solids. Culms were collected at four different time points, ranging from six to 12-month-old plants. Here we show differentially expressed genes related to sucrose metabolism and cell wall biosynthesis, including genes encoding invertases, sucrose synthase and cellulose synthase. Our results showed increased expression of invertases in IN84-58, the genotype with lower sugar and higher fiber content, as well as delayed expression of secondary cell wall-related cellulose synthase for the other genotypes. Interestingly, genes involved with hormone metabolism were differentially expressed across time points in the three genotypes with higher soluble solids content. A similar result was observed for genes controlling maturation and transition to reproductive stages, possibly a result of selection against flowering in sugarcane breeding programs. These results indicate that carbon partitioning in apical culms of contrasting genotypes is mainly associated with differential cell wall biosynthesis, and may include early modifications for subsequent sucrose accumulation. Co-expression network analysis identified transcription factors related to growth and development, showing a probable time shift for carbon partitioning occurred in 10-month-old plants.

## 1 Introduction

Brazil is the main sugarcane producer in the world, followed by India, Thailand and China (Food and Agriculture Organization, 2021). Sugarcane is the most efficient plant at accumulating sucrose (Waclawovsky et al., 2010) and is widely used as raw material for sugar and ethanol production. The bioethanol produced from sugarcane can reduce dependence on fossil fuels, which are limited resources responsible for increasing greenhouse gas emissions (Flugge et al., 2017).

Currently used commercial sugarcane genotypes are hybrids, mainly between the species *S. officinarum* and *S. spontaneum*. The former was used because of its higher sugar content in its culms, and the latter for its robustness, resistance to diseases (Campos and Caligari, 2017) and abiotic stress tolerance (Zhang et al., 2018). Following hybridization, successive backcrossing with *S. officinarum* was used to increase sugar content yield, a process called “nobilization” (Piperidis et al., 2020). These noble canes exhibit lower fiber content, thicker culms and recalcitrant flowering when compared to *S. spontaneum* (Kaffka and Grantz, 2014). Also, modern sugarcane cultivars (*Saccharum* X *officinarum*) are aneuploid and autopolyploid, with a basic ploidy level of 10x to 12x (Piperidis and D’Hont, 2020), resulting in a total genome size of roughly 10 Gbp (D’Hont, 2005).

Breeding for sugar accumulation has resulted in a sucrose content of 12-16% of the fresh weight and 50% in dry weight of mature culms (Bull and Glasziou, 1963). The accumulation of sucrose in sugarcane culms occurs by stopping the active growth and elongation in some internodes (Bindon and Botha, 2002). In mature culms, sucrose produced in mesophyll cells is unloaded to storage cells via the apoplast or symplast (Rae et al., 2005), the latter being the most frequent type of transport. This symplastic pathway can follow two routes: translocation of sucrose into the cell mediated by sucrose transporters located in the plasma membrane (Rae et al., 2005); or via sucrose hydrolyzation into fructose-6-P and UDP-glucose by acid invertases. Sucrose phosphate synthase (SPS) subsequently converts these two products into sucrose-P, which is next converted into sucrose by the activity of sucrose phosphate phosphatase (SPP) (Partida et al., 2021). A peak in sucrose levels is achieved at the end of the vegetative cycle, when this carbohydrate is used during flowering and seed production (Slewinski, 2012). For this reason, maturing and/or mature sugarcane culms are commonly used as material for the study of carbohydrate partitioning and metabolism related to sucrose accumulation (Verma et al., 2011b; Thirugnanasambandam et al., 2017). Immature culms arise from apical meristems and, in turn, are responsible for increasing the height and width of sugarcane culms by successive cell divisions (Moore and Botha, 2014), providing a backbone for the plant.

Immature (internode -1 to +3), intermediate (internode +5) and mature culms (internode +9 or more) were previously compared in terms of gene expression changes between different maturation stages (Papini-Terzi et al., 2009; Thirugnanasambandam et al., 2017). Papini-terzi et al. (2009) used cDNA microarrays to compare culms with low and high soluble solids content (°Brix), as well as mature culms vs. immature culms. Among the differentially expressed genes, the authors reported genes associated with hormone signaling, stress response, cell wall metabolism, calcium metabolism, protein kinases, protein phosphatases and transcription factors. Mature and immature culms are part of the source-sink tissues. Nevertheless, little is known about gene regulation in the latter, especially when comparing hybrids to wild-type genotypes that contrast in sugar content. A few papers report differences between these tissues and genotypes (Papini-Terzi et al., 2009; Nishiyama et al., 2014); however, they do not cover different periods of plant development. This type of study would also make it possible to investigate the interaction between genotypes and developmental period.

Gene co-expression networks can be used to provide complementary insights to differential expression analyses. These networks are based on pairwise correlations between the expression levels of genes across many samples, which are combined into a correlation matrix (Langfelder et al., 2013). From the correlations it is possible to identify expression profiles that can be related to certain biological processes (Usadel et al., 2009). In this regard, a combination of gene expression profiles and co-expression analysis can provide more information about gene expression in apical culms of contrasting genotypes of sugarcane. In this study, we aimed to investigate the transcriptional profiles of immature sugarcane culms one wild *S. spontaneum* accession and three sugarcane modern hybrids with varying contents of soluble solids.

## 2 Materials and methods

### 2.1 Plant material

We initially chose four genotypes representing different levels of soluble solids (°Brix) in a panel of *Saccharum* genotypes (Supplementary Table 1). The soluble solids content was used as a proxy for sugar content, to classify each genotype as having very high °Brix (VHB), high °Brix (HB), low °Brix (LB) or very low °Brix (VLB). We also considered the relevance of genotypes to previous genomic studies. Hybrids SP80-3280 and R570 have been widely used in genomic studies and IN84-58 is a commonly used representative of *S. spontaneum* (Partida et al., 2021). Genotype F36-819 is an important ancestor in Brazilian breeding programs, found in the pedigree of RB867515 (Barreto et al., 2019), currently one of the most grown cultivars. These genotypes are present in the Brazilian Panel of Sugarcane Genotypes (BPSG) (Barreto et al., 2019; Medeiros et al., 2020), which currently contains 254 genotypes, including representatives of wild *Saccharum* species, relevant cultivars for breeding programs and cultivars from other sugarcane producing countries. The plants in the field were grown in a randomized block design with four replicates, at the Federal University of São Carlos/Araras (22°18′41.0″S, 47°23′05.0″W, at an altitude of 611 m) – São Paulo – Brazil.

We collected immature culms from the +1 internode of three replicates of each genotype. We sampled plants at four evenly distributed time points: April (T1), June (T2), August (T3) and October (T4) of 2016, corresponding to 6-, 8-, 10- and 12-month-old first ratoon plants. Hence, our experiment corresponded to a 4 × 4 factorial design, with a total of 48 samples, which allowed the investigation of genotype by time point interaction. Tissues were immediately frozen in liquid nitrogen and stored at - 80 °C until further processing.

### 2.2 RNA extraction and sequencing

We extracted total RNA with the *RNeasy Plant Mini kit* (QIAGEN, Valencia, CA, USA) following the manufacturer’s instructions. We assessed RNA concentration and the RNA integrity number (RIN) with the Bioanalyzer 2100 (Agilent Technologies, Santa Clara, CA, USA).

Next, we generated mRNA libraries following the Illumina TruSeq Stranded mRNA kit protocol and sequenced the 48 libraries in six lanes of an Illumina HiSeq 2500 to generate paired-end reads (2 × 100 bp).

### 2.3 Quality control, de novo transcriptome assembly and functional annotation

We use the same steps for quality control as Correr et al. (2020a). The resulting high quality reads were used for *de novo* assembly using Trinity v.2.5.1 (Grabherr et al., 2011) considering the parameters as previously described by Correr et al. (2020).

We evaluated the assembly integrity by assessing N50 statistics and annotation completeness against the Viridiplantae and *Sorghum bicolor* protein database v.3.1.1 (Goodstein et al., 2012), with BUSCO v.3 (Simão et al., 2015) and Trinity script of transcripts completeness, respectively.

The functional annotation was performed by aligning the *de novo* assembly with the non-redundant (nr) NCBI database, using Blastx v.2.6.0 (Altschul et al., 1990) with an E-value cutoff of 10^−3^, with up to 20 top hits. Complementary information of InterProScan was included to assign protein functions to the annotation based on domains and protein families. We used BLAST2GO (Conesa et al., 2005) to combine the results of Blastx and InterProScan, assigning Gene Ontology (GO) terms to the *de novo* assembly.

The R570 genome (Garsmeur et al., 2018) used in the co-expression analysis was re-annotated with the same parameters used for the transcriptome annotation. This functional annotation was also complemented with the information for transcription factors (TFs) retrieved from the Plant TFDB (Jin et al., 2017) and Grassius (Yilmaz et al., 2009) databases.

### 2.4 Differential expression and functional enrichment analyses

We used Salmon v.0.13.1 (Patro et al., 2017) to quantify gene expression levels of the *de novo* assembly and the R package tximport (Soneson et al., 2015) to aggregate counts at the gene level. The differential expression analyses were performed with the EDGER package v.3.24.3 (Robinson et al., 2010). First, we used a filter of at least one count per million (CPM) in at least three samples aiming to filter out genes expressed at a low level. We then fitted the full model according to the factorial design, using as factors the °Brix group and the time point of sample collection. We initially tested for marginal °Brix group effects using the VLB group – IN84-58 – as the reference level. Next, we assessed the genotype by time interaction by comparing each time point for each genotype against the first sampling time (6-month-old) of the VLB group, applying a filter of LogCPM > 1. For all the comparisons the null hypothesis *H*_0:_ *LogFC* = 0 was tested, where *LogFC* corresponds to the logarithm of the fold change. After correcting for multiple tests using the False Discovery Rate (FDR) (Benjamini and Hochberg, 1995), genes with *p*-value less than 0.05 were deemed as significantly differentially expressed genes (DEGs).

Functional enrichment analyses were performed using DEGs from each test with the GOSEQ package v.1.42.0 (Young et al., 2010). The GO terms were considered significantly enriched when the FDR-adjusted *p*-value was less than 0.05. We validated the expression of four genes with RT-qPCR (Supplementary Data 1).

### 2.5 Construction of a Gene Co-expression Network & Identification of Highly Connected Hubs

We first used Hisat2 v.2.1.0 (Kim et al., 2015) to align the reads to the R570 genome (Garsmeur et al., 2018) and featureCounts V.1.6.3 (Liao et al., 2014) was used to quantify the expression level of each gene. We constructed three co-expression networks based on the normalized CPM of the differentially expressed genes from the comparisons of time points T2, T3 and T4 against T1. To that end we used the Weighted Correlation Network Analysis (WGCNA v.1.68) package (Langfelder and Horvath, 2008). The adjacency matrix was calculated using *β* = 15 and the Topological Overlap Matrices (TOM) were calculated based on the adjacency matrix using a signed correlation, reducing noise and spurious associations. We used signed correlations because negative and positive correlations between genes reflect different modes of regulation of gene expression. We set the minimum number of genes inside a module to 250 and merged modules with a correlation higher than 0.75. Next, we exported nodes and edges to Cytoscape v. 3.8.0 (Shannon et al., 2003). The top 30 highly connected hub nodes in the network were identified using the Maximal Click Centrality (MCC) function, available in the CytoHubba (Chin et al., 2014).

## 3 Results

### 3.1 RNA sequencing, de novo transcriptome assembly and characterization of sequencing libraries

The 48 sequencing libraries yielded a total of 3.2 billion reads, with library sizes ranging from 59 to 81 million reads. The quality processing step removed roughly 17% of the total reads (Supplementary Table 2). High-quality reads were *de novo* assembled and generated 482,103 transcripts, representing a total of 190,356 unigenes with a transcript N50 length of 1,227 bp. The majority of unigenes were represented by one or a few isoforms, with 54.08%, 18.44% and 7.89% of the unigenes with one, two or three isoforms, respectively (Supplementary Figure 1).

Comparing the 190,356 orthologous unigenes against the Viridiplantae database, we found 64.9% complete sequences (92 single-copy and 184 duplicated BUSCOs), while 29.6% of unigenes represent fragmented sequences (126 BUSCOs) and 5.5% are missing (23 BUSCOs). Also, 57.13% of the sorghum protein sequences (15,872 of 27,781) were covered by unigenes in at least 70% of their length (Supplementary Table 3). The functional annotation assigned protein names to a total of 58,328 unigenes (or 30.64% of the total).

After removing genes expressed at low levels, our gene expression matrix encompassed 89,041 unigenes, which were initially used for exploratory data analysis of the sequenced libraries. We first used a multidimensional scaling (MDS) plot to assess similarities between the samples (Supplementary Figure 2). The first MDS component separated the IN84-58 (VLB) samples from the others, while the second component mainly separated samples from F36-819 (LB) at the bottom and R570 (HB) at the top, the latter showing some overlap with SP80-3280 (VHB). It is interesting to note that 12-month-old samples were more clearly separated from the other time points for every genotype. This observed grouping structure partly reflects patterns in the content of soluble solids, because the *S. spontaneum* genotype (IN84-58) showed a substantially lower phenotypic value, while the other three groups were more similar to each other. The VHB, HB and LB representatives are hybrids generated by crosses between the species *S. officinarum* and *S. spontaneum*, each undergoing different selective pressures in breeding programs.

### 3.2 Differential gene expression and functional enrichment

We used the VLB (*S. spontaneum*) genotype as a reference against which to test for main effects of the LB, HB and VHB hybrids. In this test, 88.04% of the 89,041 expressed genes were differentially expressed in at least one group, with 44,036 commonly found among contrasts (Figure 1-A). Also, roughly 6,000 of the DEGs were commonly found in each pairwise intersection between the °Brix groups. This high number of shared DEGs is expected given that F36-819, R570 and SP80-3280 showed similar gene expression profiles when compared to IN84-58. This result may partly be explained because the VLB genotype is a wild accession, while the three remaining genotypes are hybrid sugarcane cultivars.

**Figure 1:**
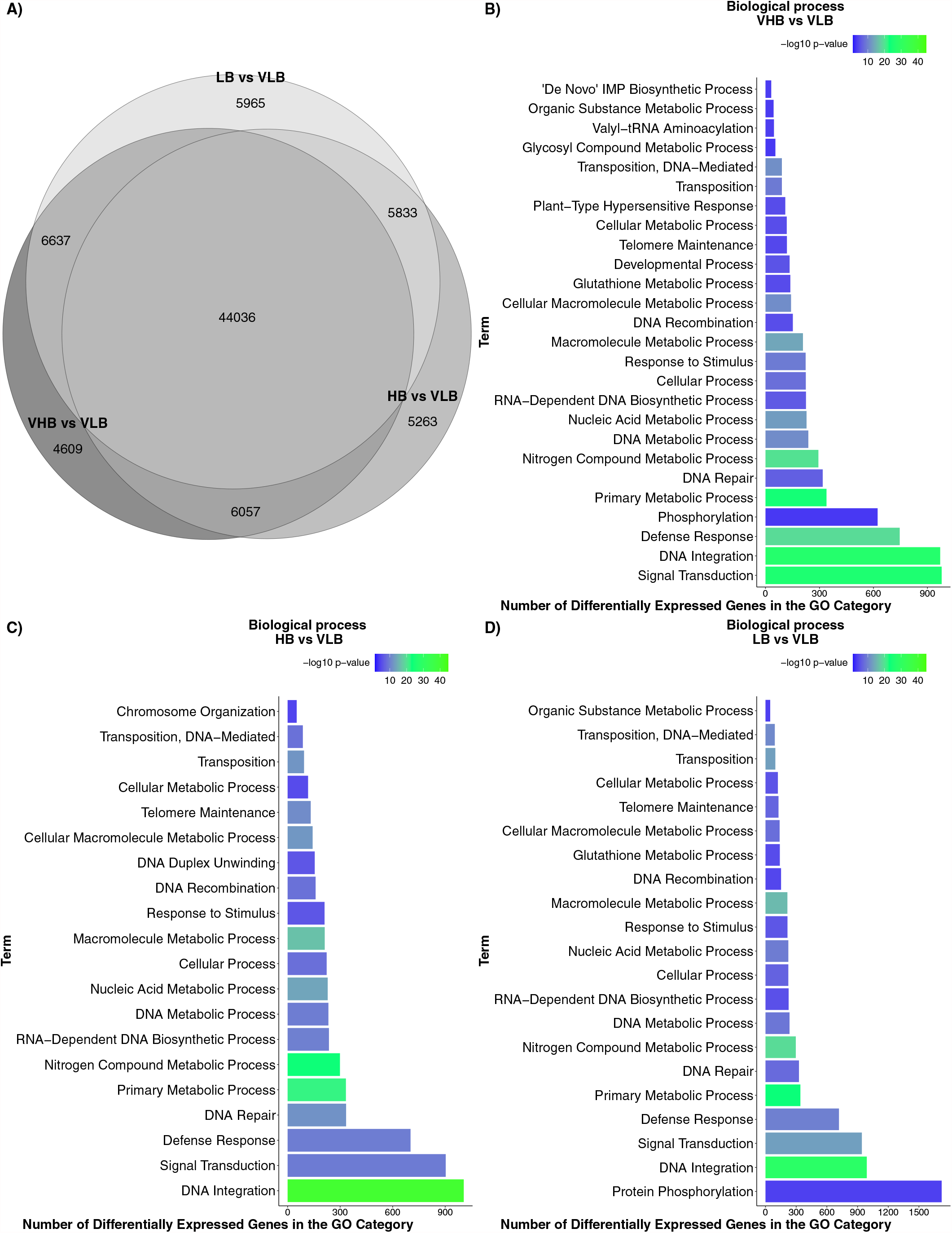
Differential gene expression and functional enrichment results: A) Venn diagram of differentially expressed genes (DEGs) found in tests for main effects. Enriched Gene Ontology (GO) terms of the biological process category: B) VHB vs VLB contrast; C) HB vs VLB contrast; and D) LB vs VLB contrast. VHB: very high °Brix, HB: high °Brix; LB: low °Brix; and VLB: very low °Brix.

Next, we identified the biological processes enriched among DEGs for each different °Brix group. We found 26, 20 and 21 enriched GO terms in the contrasts involving the VHB, HB and LB groups, respectively (Figure 1–B, C, D). Genes annotated with terms relating to transposable elements (TE), such as DNA-mediated transposition, transposition, and DNA recombination, were enriched in the three contrasts.

In all comparisons we found terms related to stress and signaling pathways, such as *defense response, response to stimulus* and *signal transduction*. Sugar accumulation or culm development can trigger the expression of genes involved with stress responses in sugarcane (Papini-Terzi et al., 2009). Also, considering that apical culms are actively growing organs, cell expansion occurs due to the reduction of osmotic potential by the accumulation of solutes. This then causes a differential gradient allowing the entry of water into the cell, resulting in an irreversible expansion of the cell (Wang and Ruan, 2013).

We performed additional tests to investigate pairwise differences between genotypes at different time points, using VLB in T1 as a reference treatment (Supplementary Figure 3). In almost all scenarios the number of downregulated genes was higher than that of upregulated DEGs. We noticed that the comparison HB vs VLB at time point T4 showed the highest number of DEGs.

### 3.3 Sucrose metabolism and transport

We investigated genes previously associated with sucrose levels and often discussed in the relevant literature. The expression profiles of nearly all invertase genes showed negative log fold changes for all combinations of genotypes and time points, compared to the reference (Figure 2-A), indicating higher expression in the VLB group than in the other three genotypes. Indeed, genes annotated as cell wall invertase (CW-INV) and soluble acid invertase (SAI) showed substantially lower expression levels from the beginning of the experiment, in most cases persisting through T3 or T4. Because these enzymes are involved in the irreversible cleavage of sucrose into glucose and fructose, this may reflect more intense breakage of disaccharides in the VLB genotype to release raw material for subsequent synthesis of cell wall components.

**Figure 2:**
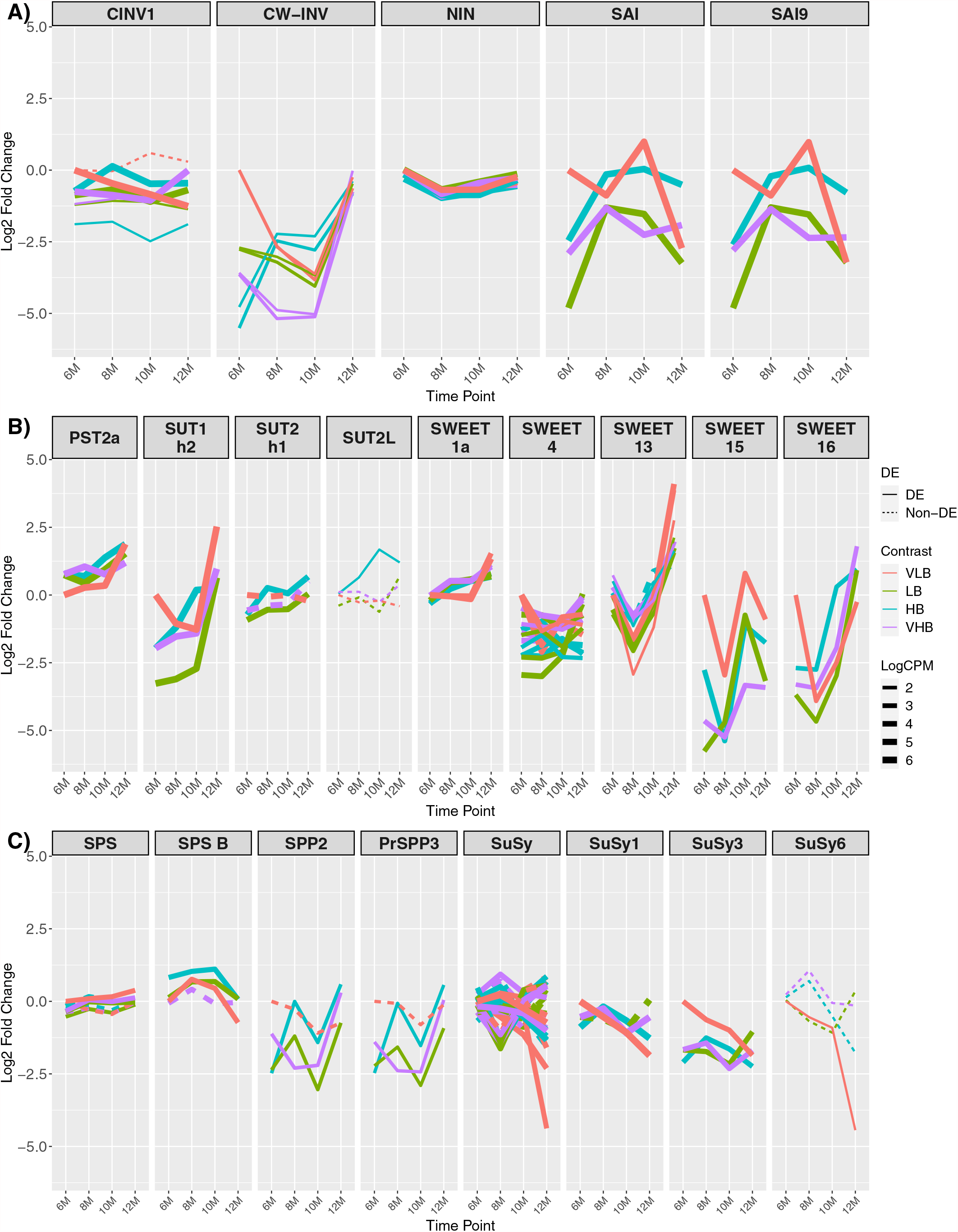
Expression profiles of genes related to sucrose metabolism. The × axis shows the age of plants (in months). The y axis corresponds to the log fold change in comparison to the very low °Brix (VLB) genotype in T1 (6-month-old plants). Each line represents a gene and the solid lines indicate genes with significant differential expression in at least at one time point, while dashed lines represent genes with no differential expression for a given genotype. The line width indicates the average expression level of each gene, with more highly expressed genes thicker. (A) Invertase genes, where CINV, CW-INV, NIN and SAI correspond to cytosolic invertase, cell wall invertase, alkaline/neutral invertase and soluble acid invertase, respectively. (B) Genes responsible for sugar transport, with PST, SUT and SWEET corresponding to putative sugar transporter, sucrose transporter and bidirectional sugar transporter SWEET, respectively. (C) Genes involved in sucrose metabolism, where PrSPP, SPP, SPS and SuSy correspond to probable sucrose phosphate phosphatase, sucrose phosphate phosphatase, sucrose phosphate synthase and sucrose synthase, respectively. VLB: very low °Brix, LB: low °Brix, HB: high °Brix and VHB: very high °Brix.

We observed a similar profile for genes that encode sugar or sucrose transporters, with the majority showing lower expression in 6-month-old hybrids when compared to VLB (Figure 2-B). We found ten sugar transporter SWEET members, with SWEET1a, 4, 13, 15 and 16 showing more marked changes over time, with mostly negative values of log fold change. SWEET13 is reported to be more expressed than SWEET4, 15 and 16 in mature culms. Also, SWEET4 and SWEET16 are suggested to be exporters of sucrose or to be involved in sugar accumulation (Hu et al., 2018). This indicates a possible source for cell wall biosynthesis or initial sugar accumulation in the apical culms.

We did not observe major differences in the expression profiles of SPS genes (Figure 2-C). However, SPS B showed positive values of log fold change for most of the treatments, albeit of small magnitude. Interestingly, sucrose phosphate phosphatase (PrSPP3/SPP2) coding genes were differentially expressed in LB, HB and VHB, showing expression profiles with reduced expression at nearly all collection time points. Sucrose Synthase (SuSy) expression profiles showed negative values of log fold change in most of the cases when compared to the reference (Figure 2-C). We observed a similar expression profile between invertases and SuSys, suggesting a possible (complementary) sucrose cleavage activity. SuSy catalyzes a reversible cleavage of sucrose into fructose and UDP-Glu, which are used for respiration and as cell wall components (Verma et al., 2011b).

Regarding the activity of trehalose-6-phosphate synthase (TPS) genes, the expression profiles of multiple members of this family showed negative values of log fold change in LB, HB and VHB, particularly from 6-to 10-month-old plants. Also, different genotypes showed strong increased expression level of TPS1, 7, 10 and 11 at T4, revealing a change in the expression pattern in the latest collection time point (Supplementary Figure 4). TPS catalyzes the first step in trehalose metabolism and is fundamental for embryo development (Eastmond et al., 2002), vegetative growth and transition to flowering in *Arabidopsis thaliana* (Van Dijken et al., 2004). Trehalose-6-phosphate (T6P) is an important signaling molecule and is associated with the protection of plants from abiotic stress (Garg et al., 2002). Also, T6P is linked to carbon partitioning (Delatte et al., 2011).

### 3.4 Cell wall synthesis and modification

We found 21 members of cellulose synthase (CesA) coding genes, seven of these – CesA3, 4, 7, 8, 10, 12 and a putative member – showing more pronounced changes across time points (Figure 3). The expression profile of CesA3 showed a positive value of log fold change for all Brix groups, while the other members of the gene family showed increasing expression levels over time points. The exception was the VLB genotype, which showed marked up or downregulation in T4 for many of these genes. Because CesAs are responsible for cellulose biosynthesis and in most cases the genotypes showed increasing expression levels across time points, this likely indicates ample synthesis of primary cell walls in dividing cells of these young sugarcane culms.

**Figure 3:**
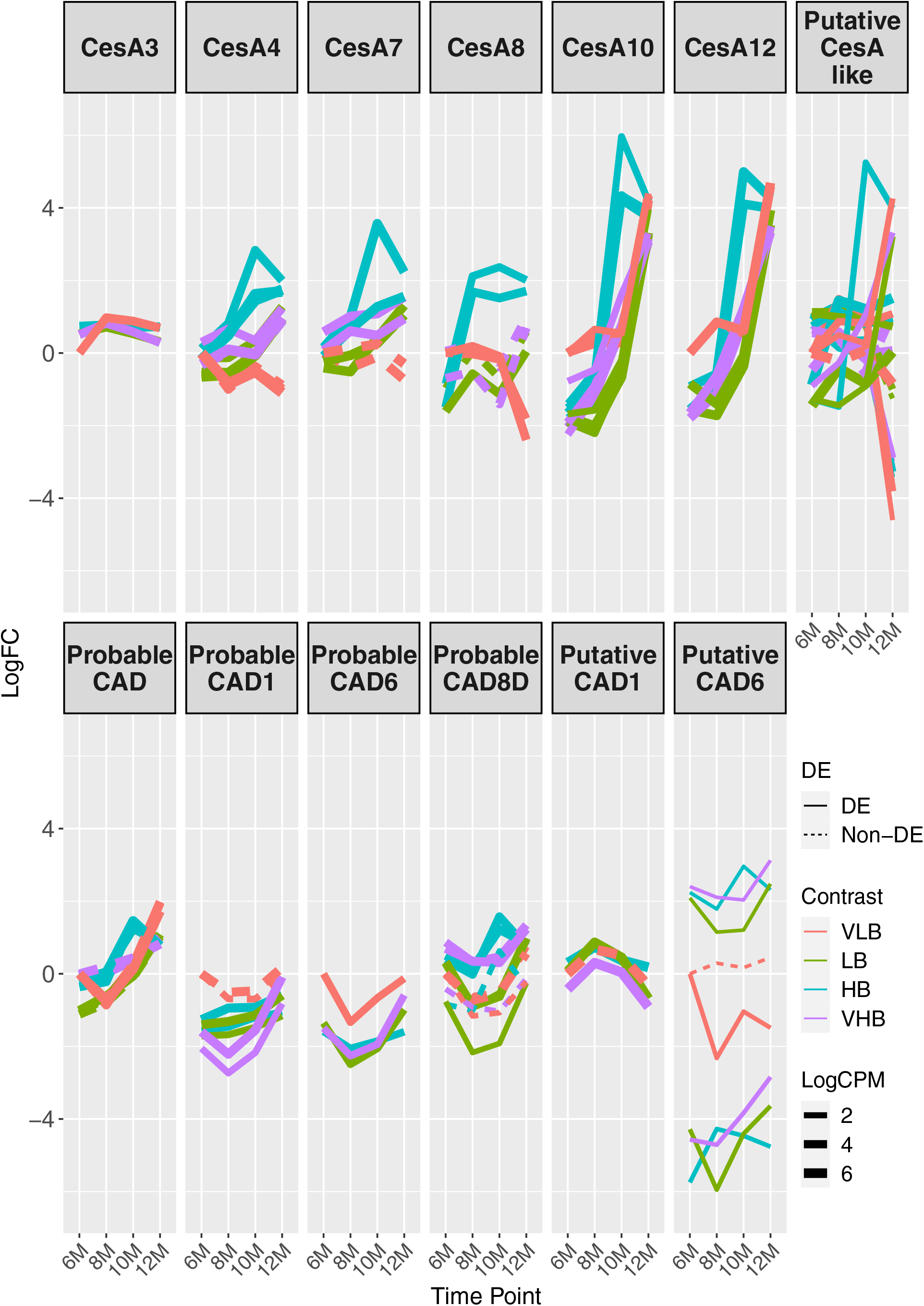
Expression profiles of genes related to the biosynthesis of cell wall components. The × axis shows the age of plants (in months). The y axis corresponds to the log fold change in comparison to the very low °Brix (VLB) genotype in T1 (6-month-old plants). Each line represents a gene and the solid lines indicate genes with significant differential expression in at least at one time point, while dashed lines represent genes with no differential expression for a given genotype. The line width indicates the average expression level of each gene, with more highly expressed genes thicker. CesA and CAD correspond to cellulose synthase and cinnamyl alcohol dehydrogenase, respectively. VLB: very low °Brix, LB: low °Brix, HB: high °Brix and VHB: very high °Brix.

Cinnamyl alcohol dehydrogenase (CAD) coding genes also revealed varying patterns of differential expression for different members of the family. The biggest changes occurred for those coding for probable CAD1 and a probable CAD6, which showed negatives values of log fold change in almost all comparisons against VLB with six months (Figure 3). CAD is a key enzyme in lignin biosynthesis (Sibout et al., 2005), and the higher expression in VLB for these young sugarcane culms shows that the differential accumulation of fiber can begin early in the development of culms. We also observed differences in the expression profiles of multiple genes encoding expansins, with many of them indicating differential expression (in both directions) in T1 and across time points (Supplementary Figure 5). This likely reflects the expansion of cells in the apical culms of all genotypes, which may depend on different members of this gene family.

### 3.5 Hormone metabolism and stress responses

We investigated gene expression profiles of genes coding for proteins associated with hormone metabolism (Figure 4) and stress responses (Figure 5) based in previous studies (Mattiello et al., 2015; Thirugnanasambandam et al., 2017). We noticed that one of the genes coding for auxin response factor (ARF23-like) and another coding for ARF75 showed strong main effects between genotypes, with increased expression level in the LB, HB and VHB groups. Conversely, one ARF6a coding gene showed consistently lower expression levels for all the genotypes, when compared to VLB. ARFs are among the transcription factors responsible for controlling auxin response genes (Guilfoyle and Hagen, 2001), which in turn control cell expansion (Mockaitis and Estelle, 2004). In addition, ARF6a is reported to regulate photosynthesis, sugar accumulation and fruit development in *Solanum lycopersicum* (Yuan et al., 2019). These observations suggest that hybrid cultivars may be preparing the apical culms for future sugar accumulation.

**Figure 4:**
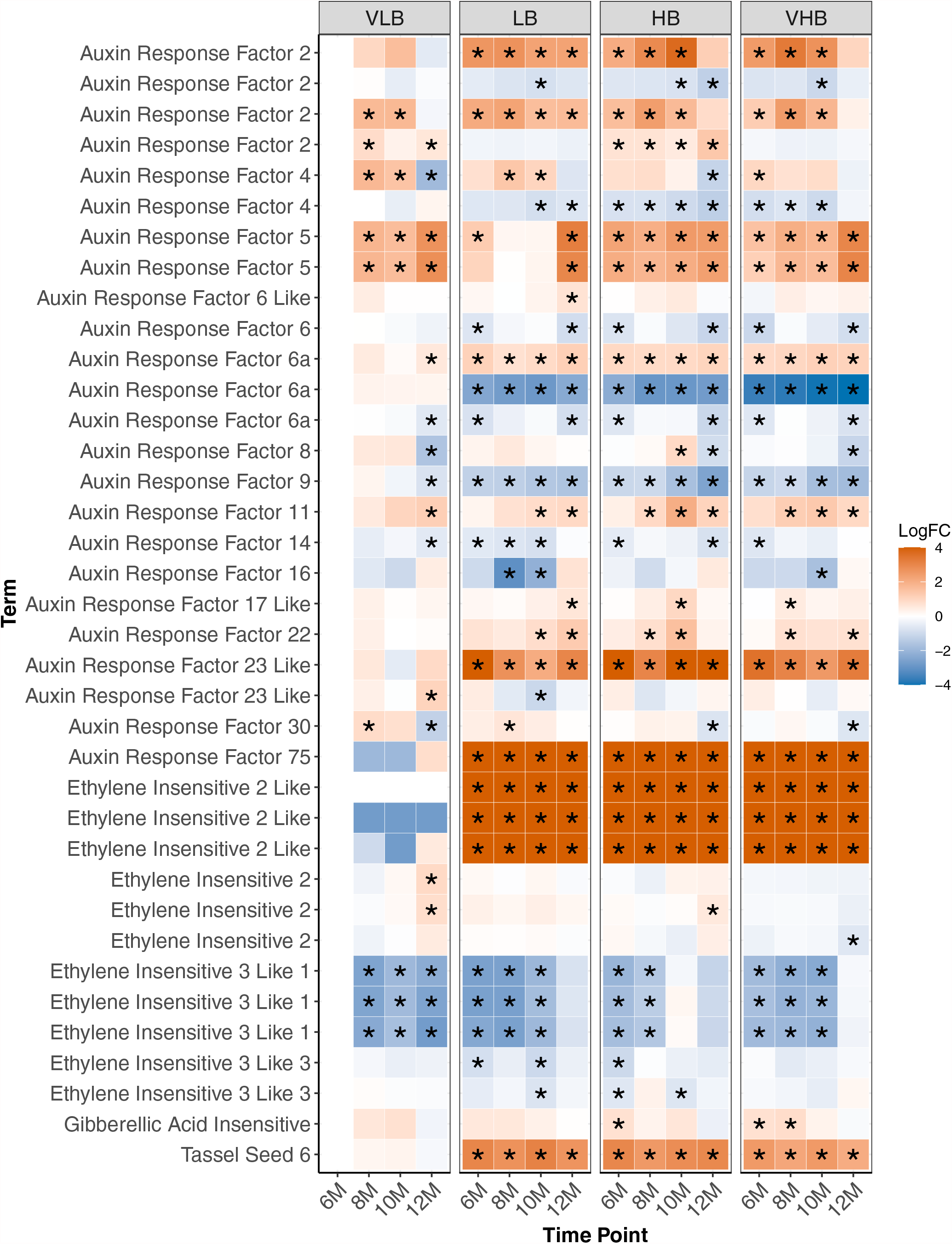
Expression profiles of genes related to hormone metabolism. Asterisks indicate genes with differential expression for a particular combination of genotype and time point. The age of plants in the × axis is shown in months. VLB: very low °Brix, LB: low °Brix, HB: high °Brix and VHB: very high °Brix. All tests considered 6-month-old VLB as a reference group.

**Figure 5:**
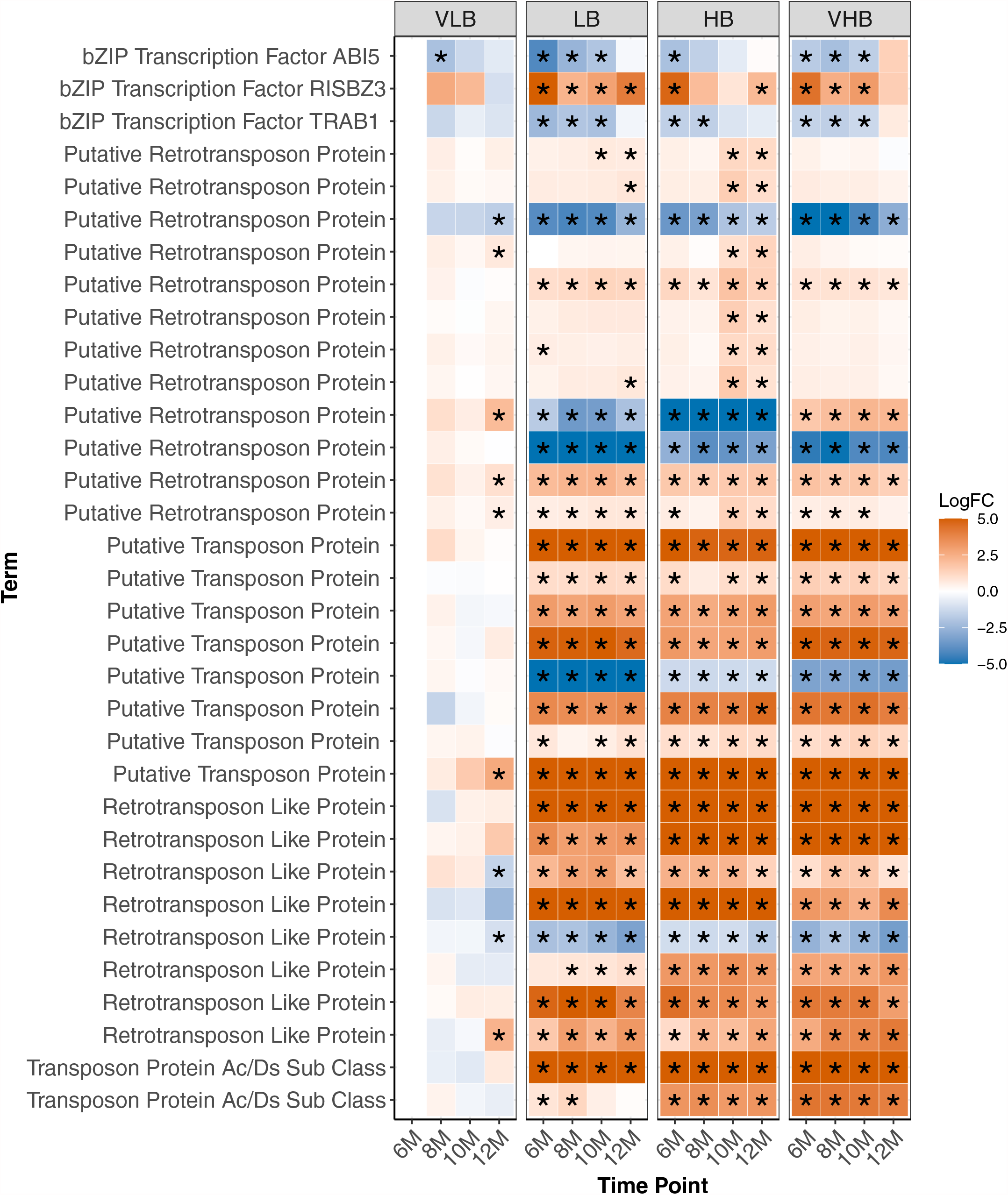
Expression profiles of genes involved in response to stress. Asterisks indicate genes with differential expression for a particular combination of genotype and time point. The age of plants in the × axis is shown in months. VLB: very low °Brix, LB: low °Brix, HB: high °Brix and VHB: very high °Brix. All tests considered 6-month-old VLB as a reference group.

Genes coding for ethylene-insensitive proteins showed varying expression patterns, with three representatives of ethylene insensitive 3 like 1 showing lower expression levels in many of the tested contrasts (Figure 4). In particular, the LB, HB and VHB genotypes showed lower expression of these genes than VLB in the first collection, but expression levels increased over time. Conversely, the VLB genotype showed decreasing expression (Figure 4). Members of ethylene insensitive 2 like (EIN2-like) showed positive values of log fold change at all time points for LB, HB and VHB, but constant expression for VLB. EIN2 rice mutants showed defective stalk development and shoot elongation (Jun et al., 2004), indicating a possible role of EIN2-like in increasing stalk size in hybrids for future sucrose accumulation. Finally, a gene coding for tassel seed 6 (TS6) also showed increased expression in the three hybrid genotypes. In maize this gene is related to sex determination and inflorescence pattern, and is also associated with delays in the development of the inflorescence meristem (Irish, 1997). The expression pattern we observed may reflect the selection against flowering of cultivars in sugarcane breeding.

We note that the majority of the genes investigated in this category showed main effects between genotypes, but little genotype × time interaction, *i*.*e*., their expression patterns were consistent across time points. Because they are mainly associated with plant development, sugar accumulation or the control of flowering, these profiles may indicate their involvement with regulating important phenotypic traits that have undergone selection after the crossing between S. *officinarum* and *S. spontaneum*.

Several transcription factors were identified, including members of the bZIP family (Supplementary Figure 6), ABI5, RISBZ3 and TRAB1. Among them, ABI5, RISBZ3 and TRAB1 showed evidence of differential expression in the initial time points for the three hybrids. The expression profile of ABI5 and TRAB1 showed reduced expression in the hybrids, while the expression of RISBZ3 was higher in these groups. In rice, ABI5 and TRAB1 are regulated by the hormone abscisic acid (ABA). ABI5 is linked to abiotic stress and plant pollen development (Zou et al., 2008), while TRAB1 controls maturation and dormancy in embryos (Hobo et al., 1999). Additionally, RISBZ3 is expressed during seed maturation (Onodera et al., 2001). The role of these transcription factors in regulating abiotic stress and maturation of meristematic tissues possibly reflects the selection performed in breeding programs to develop hybrid sugarcane genotypes.

Interestingly, most of the genes involved with stress response detected herein code for transposable elements (TEs), especially retrotransposons. Retrotransposon genes were mostly upregulated in LB, HB and VHB genotypes, with little interaction with time points (Figure 5). Similarly, we observed virtually no difference in expression levels for the four VLB collections, with a few exceptions in the more mature 12-month-old plants. Because TEs are known to introduce mutations and create plasticity in plants (Lee and Kim, 2014), the apparent overexpression in LB, HB and VHB possibly reflects the interspecific hybridization and backcrossing events during the breeding history of the crop, which potentially caused widespread genomic modifications (Ming et al., 2010).

### 3.6 Expression of transcription factors over time

In order to identify key transcription factors, we performed co-expression analyses using DEGs from pairwise comparisons between time points – 8-, 10- and 12-month-old plants, each compared against the baseline of 6-month-old samples. Highly connected genes of each module were investigated as potential factors related to sugar accumulation and the top 30 genes were selected as candidate hub genes. In the comparison involving the T2 group (Figure 6-A), we identified three TFs: B3 DNA binding domain (B3), probable WRKY transcription factor 31 (WRKY31) and ZmOrphan185.

**Figure 6:**
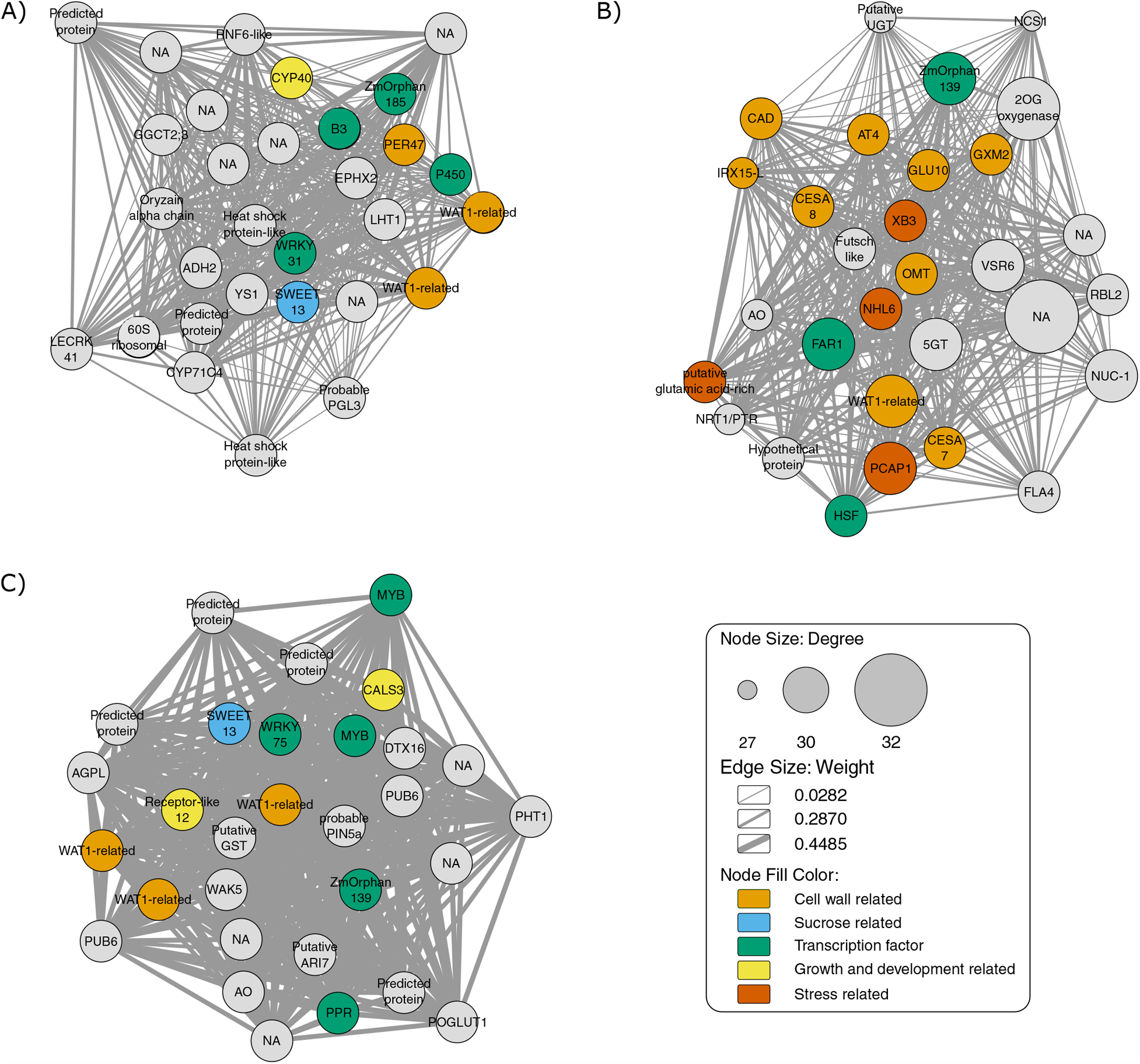
Gene co-expression network of the top 30 hub genes, as indicated by the maximal clique centrality. The node sizes represent the degree of correlation with other genes. Light orange, light blue, green, yellow, orange nodes represent cell wall related, sucrose related, transcription factor, growth and development related and stress related genes, respectively. The edges represent interactions between genes and their width indicates the corresponding weight. (A) Comparison of T2 (8-months) against T1 (6-month-old plants). (B) Comparison of T3 (10-months) against T1. (C) Comparison T4 (12-months) vs T1.

Besides these TFs, we identified the genes Walls Are Thin (related to WAT1), Peroxidase 47 (PER47) and cytochrome P450 709B2 (P450). WAT1 and PER47 are related to cell wall biosynthesis (Tokunaga et al., 2009; Ranocha et al., 2010) and P450 to salt stress in *A. thaliana* (Mao et al., 2013). WAT1-related is regulated depending on the light period and is related to growth and secondary cell wall metabolism in *A. thaliana* (Ranocha et al., 2010). In turn, PER47 is involved in vessel lignification (Tokunaga et al., 2009). Furthermore, CYP40 is necessary for the vegetative phase and floral morphogenesis of *A. thaliana* (Berardini et al., 2001). SWEET13 is involved in the process of sugar transport between source-sink tissues in the *Saccharum* genus (Hu et al., 2018). Flowering in sugarcane is a complex process and depends on the genotype, photoperiod and weather (Jadoski and Ono, 2011).

Among the TFs found in the network constructed for T3 (Figure 6-B), we found Far-Red Impaired Response1 (FAR1), HSF family protein (HSF) and ZmOrphan139. FAR1 interacts with its homolog Far-Red Elongated Hypocotyls3 (FHY3), regulating the circadian clock, flowering time (Li et al., 2011), plant defense (Wang et al., 2016), oxidative stress (Ma et al., 2016) and many other processes in *A. thaliana*. HSF also responds especially to heat stress (Nover et al., 2001). In addition, this network revealed many genes related to cell wall metabolism, including Endoglucanase 7 (GLU 10), CAD, WAT1-related, CesA7, CesA8, Glucuronoxylan 4-O-methyltransferase 2 (GXM2), Acyltransferase 4 (AT 4), Protein IRX15-LIKE (IRX15-L) and O-methyltransferase (OMT).

In this comparison, we also found genes responsive to stress such as E3 ubiquitin-protein ligase XB3 (XB3), NDR1/HIN1-like protein 6 (NHL6), plasma membrane-associated cation-binding protein 1 (PCAP1) and putative glutamic acid-rich protein. OMT participates in both process, cell wall metabolism and stress response. In A. thaliana, NHL6 is reported as being involved in the abiotic stress-induced by ABA mainly during seed germination and early seedling development (Bao et al., 2016).

Finally, the T4 comparison (Figure 6-C) showed TFs also present in the previous comparisons. We also identified genes encoding a MYB family protein (MYB) and pentatricopeptide repeat protein-like gene (PPR). The homolog of the MYB is involved in controlling biotic (Segarra et al., 2009) and abiotic stress tolerance in *A. thaliana* (Jung et al., 2008), cell shape in Antirrhinum flowers (Noda et al., 1994) and flavonoid production (Hichri et al., 2011). Another TF identified is WRKY75, which is mainly related to phosphate deprivation in *A. thaliana* (Devaiah and Raghothama, 2007). Interestingly, a post-transcriptional factor PPR was identified, and mutants of this gene in rice showed albino phenotype and abnormal chloroplasts (Gong et al., 2014), with reduction of growth and development (Sung et al., 2010). As in the previously comparisons, we identified SWEET13 and WAT1-related. Furthermore, we found other genes related to growth and development such as callose synthase 3 (CALS3) and a receptor-like protein 12. CALS3 is expressed in the stele and root meristem during *A. thaliana* development (Vatén et al., 2011), while receptor-like 12 is involved in the regulation of meristem maintenance in the same species (Wang et al., 2010).

As expected in the young apical culms, in these three co-expression networks we found TFs related to growth, development, biotic and abiotic stress responses, inflorescence development and biosynthesis of secondary metabolites. Our hypothesis is that all time points after T1 presented metabolic changes, probably as a reflection of growth and cell expansion. In addition, T3 showed more genes related to cell wall metabolism showing a possible time shift for carbon partitioning in sugarcane.

### 3.7 Sugarcane ePlant

We also have implemented a Sugarcane ePlant (http://bar.utoronto.ca/~asher/eplant_sugarcane/) at the BAR (Bio-Analytic Resource for Plant Biology). ePlant is a compilation of programs, providing a user interface framework allowing interactive visualizations at multiple scales. The data in ePlant are displayed based on a hierarchical scale, combining many tools to provide useful information (Waese et al., 2017). One of the tools available in ePlant is the *Tissue and Experiment eFP Viewer* (Figure 7), showing gene expression details on a gene-by-gene basis across the different samples in the experiment described in this paper. Here we display the visualization of the expression pattern of SWEET13 (Sh08_g006050), providing similar results as the expression profiles shown in Figure 2-B.

**Figure 7:**
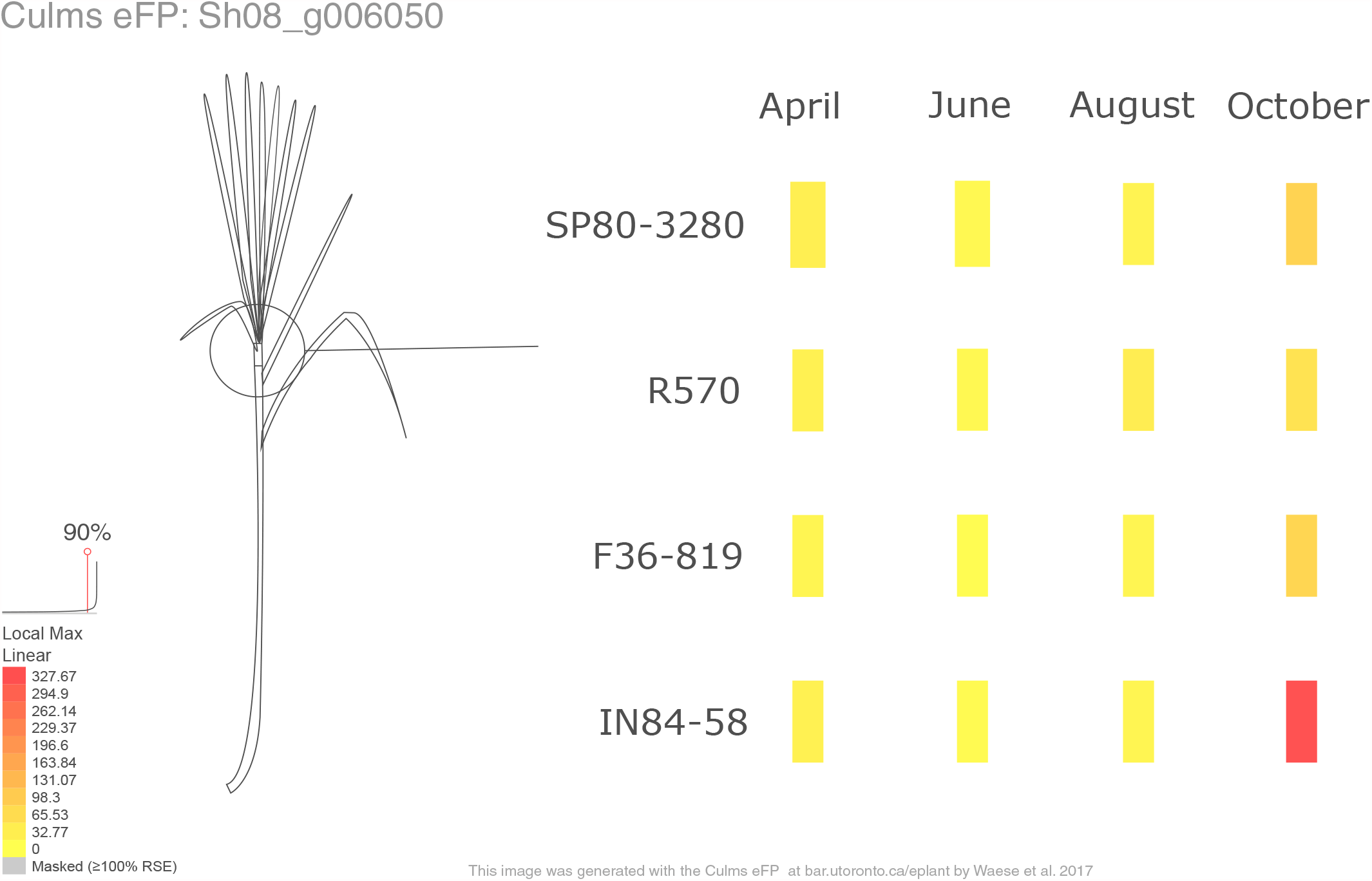
The Tissue and Experiment eFP Viewer of SWEET13 (Sh08_g006050) available in the Sugarcane ePlant. The red and yellow colors denote high and low levels of expression, respectively.

## 4 Discussion

In this study we analyzed immature culms of four sugarcane genotypes contrasting in their soluble solids content, at four time points during the growing season ranging from 6-to 12-month-old plants. Our goal was to study gene expression changes in apical culms related to carbon partitioning, aiming to understand differences between genotypes contrasting in their sugar content. The genotypes selected for this study were SP80-3280, R570, F36-819 and IN84-58, which vary regarding the soluble solids content (Supplementary Table 1). We carried out investigations during the dry season in the Center-South region of Brazil, when sugarcane normally accumulates most of its sucrose (Jadoski and Ono, 2011). We initially tested for differential gene expression considering the main effects of the genotype factor, using the very low °Brix genotype IN84-58 as a reference. We found many DEGs in common for the three genotypes with higher °Brix levels – F36-819, R570 and SP80-3280. This observation can be partially explained by breeding efforts focusing on sugar accumulation, because IN84-58 is a wild representative of the *S. spontaneum* species with higher lignin content, while all others are hybrids that have undergone selection at varying intensities. Basal metabolism, transposition and defense response were enriched among DEGs from the contrasts between groups. TEs are related to a range of mutations, mainly to substitutions, insertions and deletions of nucleotides (Kidwell, 2002). In sugarcane, TEs are widely distributed in hybrids and can introduce mutations in other coding genes, potentially resulting in greater allelic diversity (Lee and Kim, 2014). Also, TEs can be related to genome rearrangements caused by recent polyploidy (Premachandran et al., 2011). Similarly, TEs have been reported to be active under conditions such as pathogen attacks, physical injuries or abiotic stress (Grandbastien, 1998). Cell expansion occurs by the accumulation of sugar, iron or other types of solutes, creating an osmotic potential that allows water to enter into the cell, resulting in an irreversible expansion (Wang and Ruan, 2013). As a consequence, plants can trigger defense and stress response to protect from osmotic regulation (Couée et al., 2006).

To account for changes in gene expression across time points, we conducted pairwise tests for each genotype and time point combination, using as reference the expression level of VLB in 6-month-old plants. Despite the varying soluble solids content in the genotypes used in this work, the total number of DEGs was similar in all comparisons. Interestingly, differences were more marked in plants with 12 months of age (T4), probably because the hybrids were more mature than the *S. spontaneum* at this point in time.

We assessed in more detail the expression profiles of genes annotated with categories related to sucrose metabolism, cell wall biosynthesis, hormone metabolism and stress responses. Sugar accumulation is one of the best-studied processes in sugarcane, with genes reported to show up or downregulation depending on the collected material and maturation stage (Papini-Terzi et al., 2009; Thirugnanasambandam et al., 2017). Sucrose accumulation is related both to sugarcane maturation stage and internode position in the culm (Botha and Black, 2000). The sucrose produced in the leaves is transported to the culms and can be hydrolyzed by three invertases: cell-wall invertases, neutral invertases (located in the cytoplasm) and soluble acid invertases (located in the tonoplast). Invertases are key enzymes responsible for breaking sucrose into hexoses for translocation into the cell/vacuole (Ansari et al., 2013). In sugarcane, acid invertases cleave sucrose into fructose and glucose, which are subsequently translocated into the cell by transporters. Then, these monosaccharides can be used as osmotic regulators for cell expansion and growth processes (Zhu et al., 1997). In our work, cell wall invertase and soluble acid invertases showed downregulation in T1 for the three hybrids. Given that the VLB group is represented by a *S. spontaneum* accession, which accumulates more fiber than the hybrids, this suggests that differential cleavage of sucrose for deposition of cell wall components occurs in apical culms. SAIs have been previously inversely related to sugar accumulation, besides the activity of SAI decrease during the developmental stages (Verma et al., 2011a).

Another essential process for sucrose accumulation is sucrose translocation, which can occur via the apoplast or symplast. One type of symplastic transporter protein is called bidirectional sugar transporter, or SWEET (Sugars Will Eventually be Exported Transporter). SWEET13a, SWEET13b and SWEET16a are possibly associated with sugar content in species of the genus *Saccharum* (Hu et al., 2018). Our results indicate that the gene encoding SWEET13 showed increased expression levels at 12 months, suggesting an increase in the transport of photosynthates in the late stages of sugarcane development. SWEET13 was also present among the most highly connected genes in the T2 and T4 co-expression networks, curiously with contrasting gene profiles (downregulation in T2 and increased expression level in T4). These results may reflect an intensification of vegetative growth in apical culms, because of changes in the environmental conditions (Supplementary Figure 7).

Sucrose is one of the factors that contribute to plant development controlled by source-sink relations, regulating carbon translocation through the phloem and subsequent fixation in different plant organs (Lemoine et al., 2013). Sucrose accumulation is mainly correlated with the activity of SPS, followed by a sequential action of SPP (Botha and Black, 2000). The DEGs coding for SPS B showed increased expression until the last sampled time point, when its expression returned to the basal level for most of the genotypes or was down-regulated in the case of VLB. The genes encoding SPP (PrSPP3 and SPP2) showed a similar profile between comparisons, with higher expression in VLB in the initial time points. In the literature, SPS has been reported to be upregulated in mature sugarcane culms (Verma et al., 2011b; Wang et al., 2019). Our results revealed downregulation in the expression of SPP in apical culms at almost all time points. This result is in agreement with Partida et al. (2021), who reported downregulation of SPP 2D over the period of three to 12 months in source tissues of sugarcane. In any case, we did observe differences in temporal expression, which likely can be associated with the maturation of the genotypes in T4.

In plants, sucrose is converted to UDP-glucose (UDPG) regulated by the activity of SuSy, which causes a reversible cleavage of sucrose in sink tissues (Verma et al., 2011b). Also, sucrose storage in developing culms coincides with the reduction of hexoses (Whittaker and Botha, 1997). SuSy leads to the formation of UDPG and other nucleotide sugars, which can be used to synthesize cell wall components (Sung et al., 1989). In sugarcane, SuSy cleaves sucrose into UDPG for the formation of cell wall material and starch (Lingle and Smith, 1991). In addition, SuSy is associated with internode elongation (Lingle and Smith, 1991) and may be related to sucrose synthesis in immature internodes (Goldner et al., 1991). SuSy1 and SuSy3 showed higher expression in VLB in 6-month-old plants (T1), but after eight months (T2) SuSy1 was more expressed in the hybrids. These hybrids did not show differential expression of SuSy6, while the *S. spontaneum* genotype showed reduced expression over time. A previous study showed a decrease in the activity of SuSy in immature culms and mature culms of different genotypes with low and high sugar levels across time, however with less intensity in high sugar levels (Verma et al., 2011b). This observation is probably because younger culms require more hexose sugars for cell wall formation and growth processes, with demand decreasing during maturation.

Cell wall biosynthesis can be negatively correlated with sucrose accumulation, by changing the carbon partitioning towards cell wall expansion (Casu et al., 2015). Cellulose is a major component of plant biomass and is responsible for giving rigidity to the cell wall, providing the backbone to determine cell shape and maintaining the resistance of turgor pressure (Gu and Somerville, 2010). Cellulose biosynthesis in different species depends on specific members of the CesA family. In *A. thaliana* CesA4 is related to the deposition of the secondary cell wall (Taylor et al., 2003), while CesA10 and CesA12 are responsible for the same process in *Zea mays* (Appenzeller et al., 2004) and CesA7 and CesA8 in *Oryza sativa*. Besides that, CesA3 is related to primary cell wall biosynthesis in maize (Appenzeller et al., 2004). CesA10 and CesA12 showed lower expression in LB, HB and VHB groups in T1. For HB the expression was upregulated in T3, while for VLB, LB and VHB it was upregulated in T4. The expression levels of CesA4 and CesA7 increased over time for the LB, HB and VHB groups, and decreased for VLB. CesA8 showed variable profiles between soluble solids groups. The downregulation of CesA4 and CesA7 in VLB across time points was in accordance with the results obtained in immature culms of high fiber sugarcane genotypes (Kasirajan et al., 2018). Besides that, Casu *et al*. (2015) found upregulation of CesA10, 11, 12 in the rind, in comparison to vascular parenchyma and vascular bundles of mature sugarcane internodes. Primary and secondary cell wall biosynthesis is a dynamic process and is active in immature and mature culms of sugarcane (Casu et al., 2007). In our case, CesA10 and CesA12 were upregulated in T1 of VLB, probably because VLB represents a wild and more fibrous accession. In addition, CesA7 and CesA8 were respectively up and downregulated in T3 (ten months), showing high correlation in the co-expression network for this time point, with high connection to cell wall metabolism. This observation suggests a time shift in carbon partitioning in the apical culms.

Lignin is an organic polymer that is one of the secondary cell wall components, being essential for the cell wall backbone and rigidity (Chabannes et al., 2001). The final reduction of alcohols in the monolignols/lignin biosynthesis pathway is catalyzed by CAD (Sibout et al., 2005). This process can be regulated depending on the maturation phase (Sauter and Kende, 1992) or by abiotic/biotic stress in monocots (Mitchell et al., 1994). In *O. sativa* CAD1-9 are related to monolignols/lignin biosynthesis (Tobias and Chow, 2005), and CAD members C and D are related to lignin biosynthesis to sustain the floral stem development in *A. thaliana* (Sibout et al., 2005). We noticed the increased expression level of genes coding probable CAD1 and probable CAD6 in the VLB group. These results agree with higher lignification of the immature tissues during the early developmental stages of the high fiber genotype IN84-58. Conversely, Vicentini et al. (2015) reported low expression levels of CAD6 in mature sugarcane culms of two genotypes contrasting in lignin content.

Plant hormones are crucial for plant development and signaling networks, as well as responses to stresses (Bari and Jones, 2009). Auxin is a key hormone for plant growth regulating tropisms, apical dominance, root development, cell expansion, division and differentiation (Hagen and Guilfoyle, 2002). ARFs are exclusive to plants and correspond to a family of TFs that control the expression of auxin response genes (Guilfoyle and Hagen, 2007). ARFs are also related to plant development and stress tolerance (Van Ha et al., 2013). In our results, one ARF23-like and ARF75 coding genes showed differences between groups with varying contents of soluble solids, with increased expression in LB, HB and VHB. The phytohormone ethylene is related to many developmental and growth processes and stress signaling. The corresponding synthetic hormone (ethephon) is reported to be related to sucrose accumulation in immature sugarcane stems by affecting internode elongation (Cunha et al., 2017). EIN2 plays a central part in controling the ethylene signaling pathway in *A. thaliana* (Ju and Chang, 2015). In our work, EIN2-like and TS6 showed increased expression levels in the hybrids, when compared to the wild *S. spontaneum* accession. Mutations in TS6 in maize are related to sex determination and inflorescence branching, and this gene is associated with delays in the development of the inflorescence meristem (Irish, 1997). In our experiment ARF23-like, ARF75, EIN2-like and TS6 were upregulated in LB, HB and VHB, which are hybrids from sugarcane breeding programs. This suggests that *S. officinarum* may have contributed to the recalcitrance of flowering in the “nobilization” process when compared to *S. spontaneum*. (Kaffka and Grantz, 2014).

The concept of physiological stress is related to how the plant responds to many environmental conditions that limit growth, development, reproduction and crop productivity (Moore and Botha, 2014). bZIP TFs control processes related to pathogen attack (Droge-Laser et al., 1997), light and stress signaling (Choi et al., 2000), seed maturation and flowering development (Xiang et al., 1997). In rice, bZIP ABI5 and TRAB1 are regulated by ABA. Zou et al. (2008) described that overexpression of ABI5 resulted in plants with increased sensibility to salt stress, while repression lead to plants with higher tolerance but with abnormal pollen development, TRAB1 is described to control maturation and dormancy in embryos (Hobo et al., 1999). Also, bZIP RISBZ3 is expressed in the late stages of maturing seeds in rice (Onodera et al., 2001). In our results, the bZIP TFs ABI5 and TRAB1 showed downregulation, whereas RISBZ3 showed increased expression level at the initial time points for the LB, HB and VHB groups. This suggests a possible role in controlling the abiotic stress caused by the development of young tissues, as well as in the early control of maturation in young sugarcane culms (Moore and Cosgrove, 1991). This role is possibly shared with P450, FAR1, WRKY and MYB, TFs found in the temporal co-expression networks. All these TFs are mainly related to growth and development or stress responses, suggesting a sugarcane response to stress during the cell expansion process, as also reported by Carson & Botha (2002).

Transposons and retrotransposons are classes of transposable elements responsible for genome maintenance and diversification in sugarcane (Premachandran et al., 2011). Genes annotated as probable transposons and retrotransposons were mostly upregulated in all hybrids. Sugarcane hybrids represent a complex polyploid system formed by homoeologous chromosomes of *S. officinarum* and *S. spontaneum*. This possibly triggers genomic stress in these plants, where many transposable elements are activated in response to the combination of chromosomes of different species (Premachandran et al., 2011).

The accumulation of sucrose in the culms of sugarcane is controlled by a complex network of genes involved in the hydrolysis, re-synthesis and transport of sucrose, cell wall biosynthesis and abiotic stress responses. Here we found that invertases, SWEET13, SuSy1, SuSy3, probable CAD1 and CAD6 are likely related to early processes associated with carbon partitioning in apical culms. Moreover, we found evidence of particular TFs as important switches in processes related to sugarcane growth and development in early stages of maturation, as well as a probable time shift for carbon partitioning at ten months. Additionally, the TFs ARF23-like, ARF25, EIN2-like and TS6 were upregulated only in hybrids, a possible reflection of selection in sugarcane breeding programs. Although multiple studies have been developed in sugarcane, there are still many gaps in completely understanding how it accumulates sucrose in its culms. In this regard, our work provides useful gene expression profiles in apical culms of different genotypes, which can be used as genetic biomarkers for further studies.

## Supplementary data

**Supplementary Figure 1:** Distribution of the number of isoforms per unigene in the de novo assembly of the sugarcane transcriptome.

**Supplementary Figure 2**. Multidimensional scaling (MDS) plot based on gene expression profiles of immature sugarcane culms. The figure shows grouping of samples from the same genotype, with prominent separation of IN84-58 samples (very low °Brix) from the rest. Samples from 12-month-old (T4) are more clearly separated from the other time points. VLB: very low °Brix, LB: low °Brix, HB: high °Brix and VHB: very high °Brix. T1: 6-month-old, T2: 8-month-old, T3: 10-month-old and T4: 12-month-old.

**Supplementary Figure 3**. Numbers of differentially expressed genes (DEGs) per genotype. The × axis shows the age of plants (in months). Upregulated or downregulated genes are presented separately. VLB: very low °Brix, LB: low °Brix, HB: high °Brix, VHB: very high °Brix. Treatment group VLB at T1 was used as a reference for pairwise tests involving combinations of genotype and time points.

**Supplementary Figure 4**. Expression profiles of genes encoding for trehalose-6-phosphate synthase. The × axis shows the age of plants (in months). The y axis corresponds to the log fold change in comparison to the very low °Brix (VLB) genotype in T1 (6-month-old plants). Each line represents a gene and the solid lines indicate genes with significant differential expression in at least at one time point, while dashed lines represent genes with no differential expression for a given genotype. The line width indicates the average expression level of each gene, with more highly expressed genes thicker. TPS: trehalose-6-phosphate synthase. VHB: very high °Brix, HB: high °Brix; LB: low °Brix; and VLB: very low °Brix.

**Supplementary Figure 5**. Expression profiles of genes of the expansin family. The × axis shows the age of plants (in months). The y axis corresponds to the log fold change in comparison to the very low °Brix (VLB) genotype in T1 (6-month-old plants). Each line represents a gene and the solid lines indicate genes with significant differential expression in at least at one time point, while dashed lines represent genes with no differential expression for a given genotype. The line width indicates the average expression level of each gene, with more highly expressed genes thicker. EXP: expansin. VHB: very high °Brix, HB: high °Brix; LB: low °Brix; and VLB: very low °Brix.

**Supplementary Figure 6**. Expression profiles of transcription factors of the bZIP family. Asterisks indicate genes with differential expression for a particular combination of genotype and time point. The age of plants in the x axis is shown in months. VLB: very low °Brix, LB: low °Brix, HB: high °Brix and VHB: very high °Brix. All tests considered 6-month-old VLB as a reference group.

**Supplementary Figure 7**. Seasonal variation in the field in the period from Abril to October of 2016 obtained from the Automatic Weather Station (AWS) at UFSCar, Araras, SP, Brazil. A) Average temperature (°C). B) Average relative air humidity (%). C) Global solar radiation (MJ/m2). D) Total rain (mm).

**Supplementary Table 1**: Soluble solids content, type and classification of each sugarcane genotype used in this study. Plants were evaluated with a digital refractometer to estimate the content of soluble solids (°Brix). The type column indicates the genomic background of each accession, while the classification column indicates the factoring used for differential expression tests.

**Supplementary Table 2**. Number of sequencing reads before and after processing, for each sample for each Brix group.

**Supplementary Table 3**: Coverage of Sorghum bicolor proteins by the de novo assembled sugarcane transcriptome. Sorghum proteins are binned according to the percentage of coverage by at least one sugarcane transcript.

**Supplementary Data 1**. Validation of the RNA-Seq expression analysis with RT-qPCR

## 6 Author Contributions

GRAM, MSC and APS conceived and designed the experiments. MSC provided the sugarcane biological material. GRAM, GKH, FHC, FZB, and TWAB performed the experiments and data collection. GKH, FHC, CCS and DAS analyzed the data. GKH, AP and NJP designed the Sugarcane ePlant. GKH wrote the first draft of the manuscript, and all authors commented on previous versions of the manuscript. All authors read and approved the final manuscript.

## 7 Acknowledgments

This research was supported by grant #2015/22993–7, São Paulo Research Foundation (FAPESP), awarded to GRAM. This study was also financed in part by the Coordenação de Aperfeiçoamento de Pessoal de Nível Superior - Brasil (CAPES) - Finance Code 001. GKH received a fellowship from the Coordenação de Aperfeiçoamento de Pessoal de Nível Superior (CAPES - Biologia Computacional - 88882.160212/2017-01/PrInt - 88887.371113/2019-00).

## 8 Data Availability

The datasets PRJEB40481 for this study can be found in the DDBJ/EMBL/GenBank [https://www.ebi.ac.uk/ena/browser/view/PRJEB40481].

