## Supplementary Figure 1 for "Temporal Gene Expression in Apical Culms Shows Early Changes in Cell Wall Biosynthesis Genes in Sugarcane"

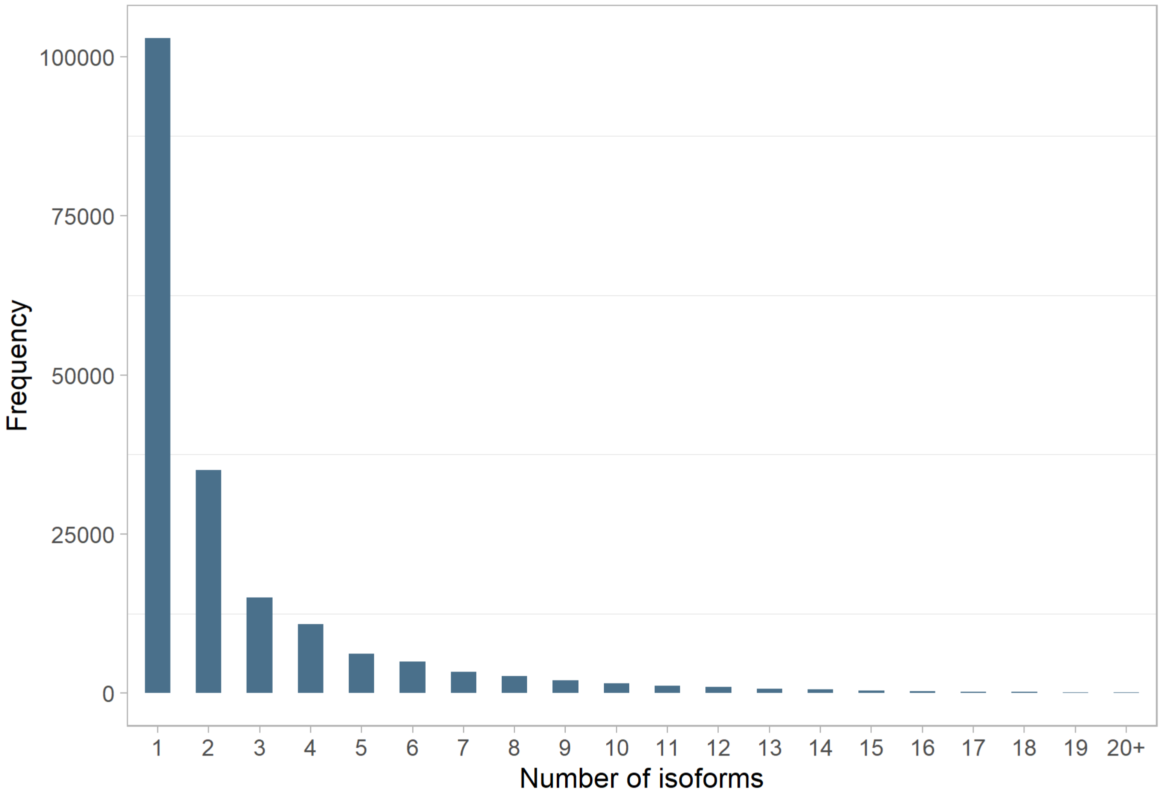


**Supplementary Figure 1.** Distribution of the number of isoforms per unigene in the de novo assembly of the sugarcane transcriptome.
