## Supplementary Figure 2 for "Temporal Gene Expression in Apical Culms Shows Early Changes in Cell Wall Biosynthesis Genes in Sugarcane"

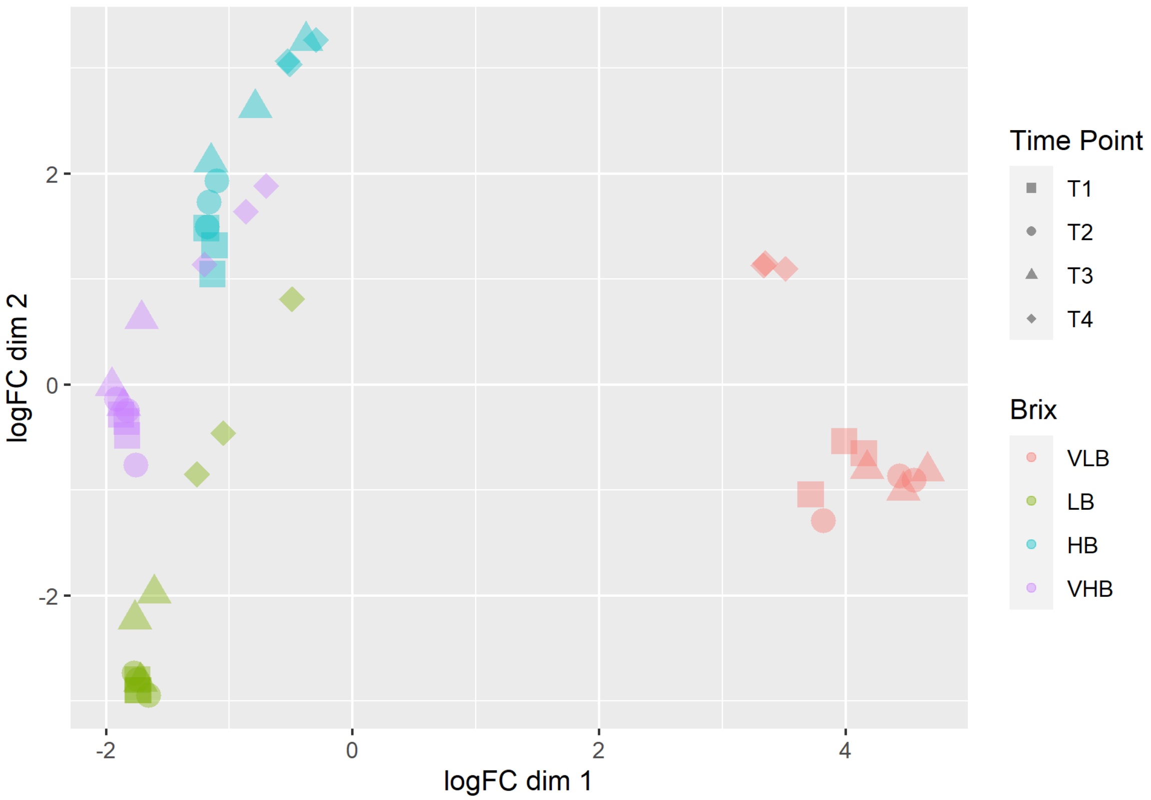


**Supplementary Figure 2.** Multidimensional scaling (MDS) plot based on gene expression profiles of immature sugarcane culms. The figure shows grouping of samples from the same genotype, with prominent separation of IN84-58 samples (very low ºBrix) from the rest. Samples from 12-month-old (T4) are more clearly separated from the other time points. VLB: very low ºBrix, LB: low ºBrix, HB: high ºBrix and VHB: very high ºBrix. T1: 6-month-old, T2: 8-month-old, T3: 10-month-old and T4: 12-month-old.
