## Supplementary Figure 3 for "Temporal Gene Expression in Apical Culms Shows Early Changes in Cell Wall Biosynthesis Genes in Sugarcane"

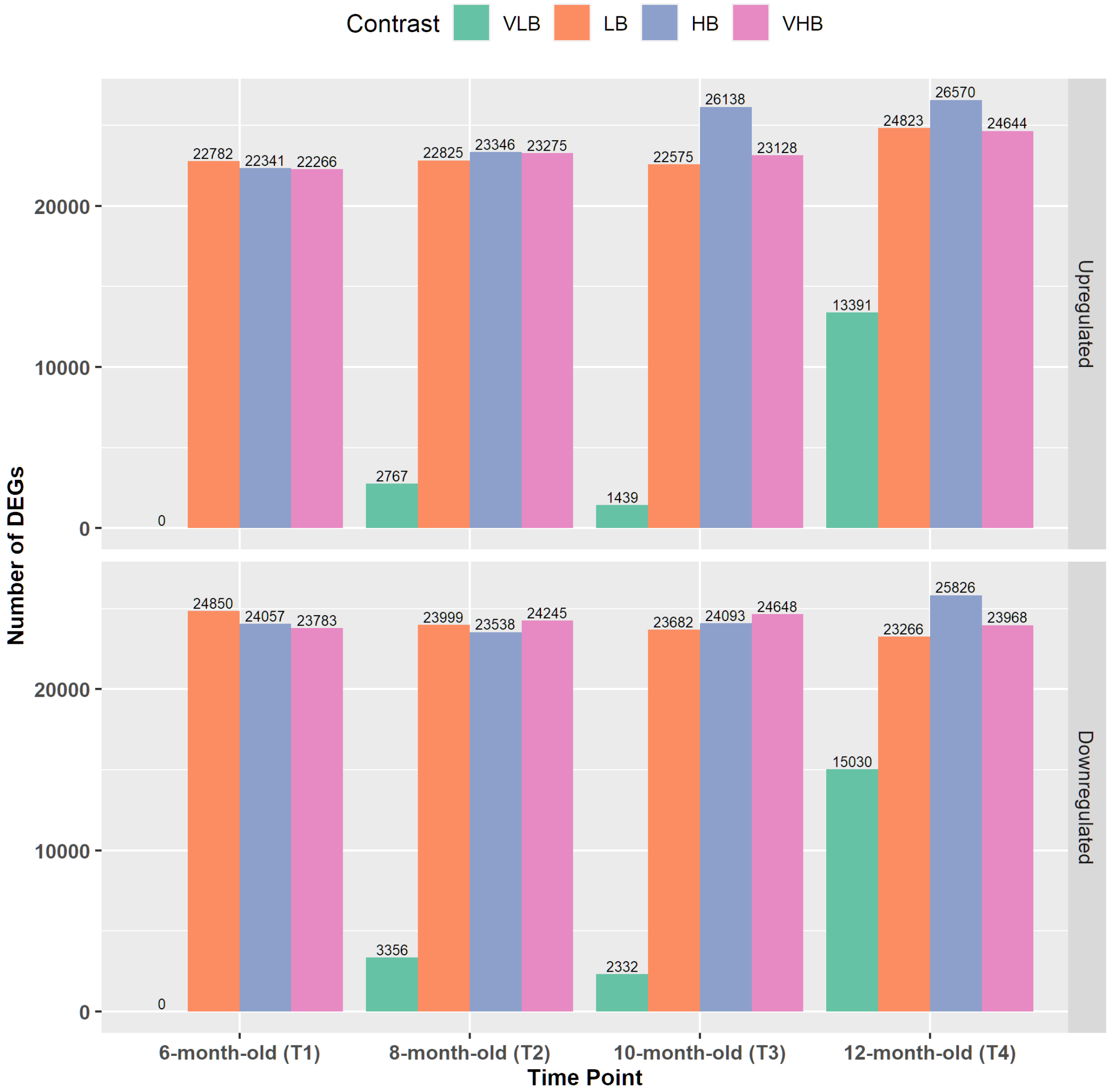


**Supplementary Figure 3.** Numbers of differentially expressed genes (DEGs) per genotype. The x axis shows the age of plants (in months). Upregulated or downregulated genes are presented separately. VLB: very low ºBrix, LB: low ºBrix, HB: high ºBrix, VHB: very high ºBrix. Treatment group VLB at T1 was used as a reference for pairwise tests involving combinations of genotype and time points.
