## Supplementary Figure 5 for "Temporal Gene Expression in Apical Culms Shows Early Changes in Cell Wall Biosynthesis Genes in Sugarcane"

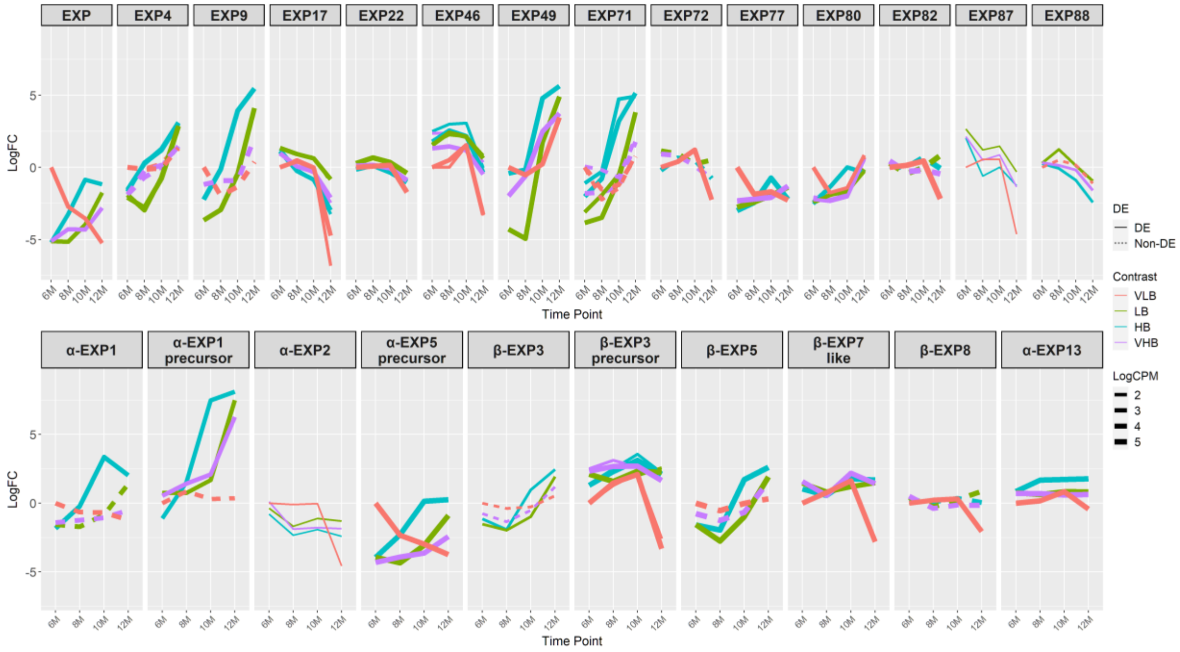


**Supplementary Figure 5.** Expression profiles of genes of the expansin family. The x axis shows the age of plants (in months). The y axis corresponds to the log fold change in comparison to the very low ºBrix (VLB) genotype in T1 (6-month-old plants). Each line represents a gene and the solid lines indicate genes with significant differential expression in at least at one time point, while dashed lines represent genes with no differential expression for a given genotype. The line width indicates the average expression level of each gene, with more highly expressed genes thicker. EXP: expansin. VHB: very high ºBrix, HB: high ºBrix; LB: low ºBrix; and VLB: very low ºBrix.
