## Supplementary Figure 6 for "Temporal Gene Expression in Apical Culms Shows Early Changes in Cell Wall Biosynthesis Genes in Sugarcane"

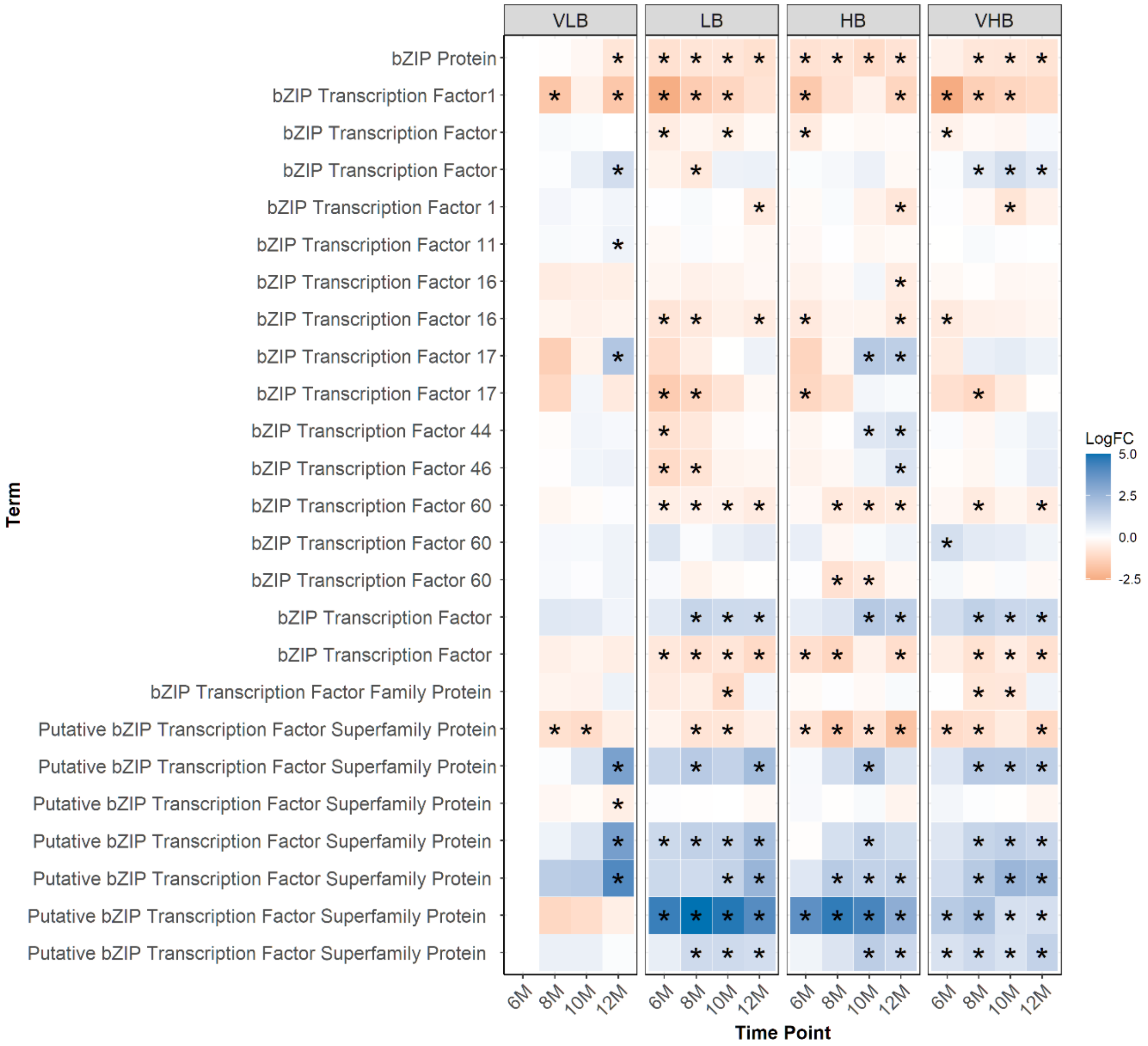


**Supplementary Figure 6.** Expression profiles of transcription factors of the bZIP family. Asterisks indicate genes with differential expression for a particular combination of genotype and time point. The age of plants in the x axis is shown in months. VLB: very low ºBrix, LB: low ºBrix, HB: high ºBrix and VHB: very high ºBrix. All tests considered 6-month-old VLB as a reference group.
