## Supplementary Figure 7 for "Temporal Gene Expression in Apical Culms Shows Early Changes in Cell Wall Biosynthesis Genes in Sugarcane"

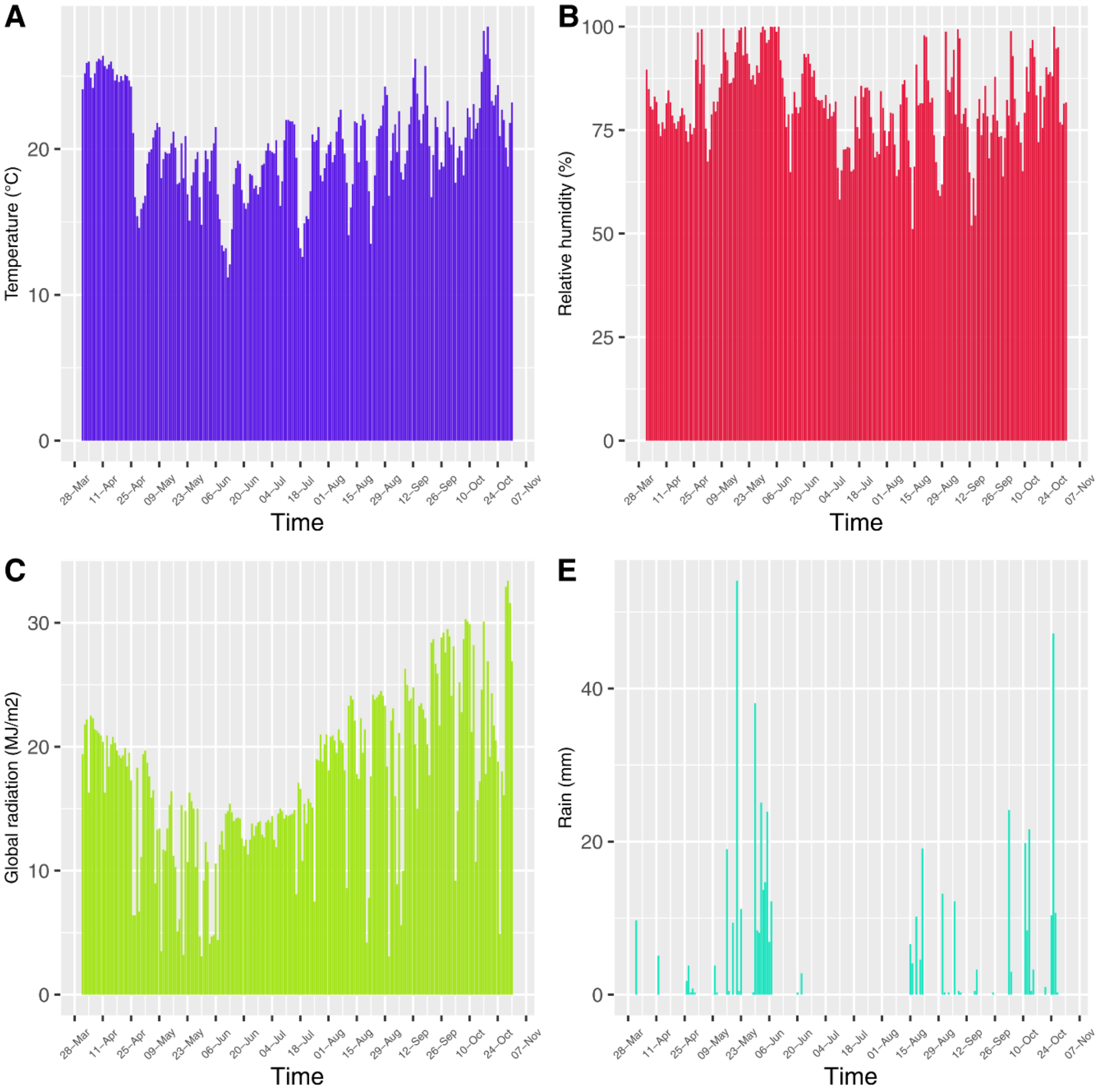


**Supplementary Figure 7.** Seasonal variation in the field in the period from Abril to October of 2016 obtained from the Automatic Weather Station (AWS) at UFSCar, Araras, SP, Brazil. A) Average temperature (℃). B) Average relative air humidity (%). C) Global solar radiation (MJ/m2). D) Total rain (mm).
