## Supplementary Table 1 for "Temporal Gene Expression in Apical Culms Shows Early Changes in Cell Wall Biosynthesis Genes in Sugarcane"

**Supplementary Table 1**: Soluble solids content, type and classification of each sugarcane genotype used in this study. Plants were evaluated with a digital refractometer to estimate the content of soluble solids (ºBrix). The type column indicates the genomic background of each accession, while the classification column indicates the factoring used for differential expression tests.

| Genotype | ºBrix | Type | Classification |
|---|---|---|---|
| SP80-3280 | 21.29 | Hybrid | Very High Brix (VHB) |
| R570 | 20.69 | Hybrid | High Brix (HB) |
| F36-819 | 18.05 | Hybrid | Low Brix (LB) |
| IN84-58 | 14.78 | *Saccharum spontaneum* | Very Low Brix (VLB) |
