## Supplementary Table 2 for "Temporal Gene Expression in Apical Culms Shows Early Changes in Cell Wall Biosynthesis Genes in Sugarcane"

**Supplementary Table 2**. Number of sequencing reads before and after processing, for each sample for each Brix group.

| Brix Group | Time Point | Replicate | Number of raw reads | Number of reads after processing | Remaining reads (%) |
|---|---|---|---|---|---|
| VLB | T1 | 1 | 68,385,684 | 56,277,658 | 82.29 |
| | | 2 | 67,523,988 | 56,092,008 | 83.07 |
| | | 3 | 72,548,456 | 60,414,788 | 83.28 |
| | T2 | 1 | 75,502,286 | 62,935,180 | 83.36 |
| | | 2 | 76,766,238 | 64,951,184 | 84.61 |
| | | 3 | 66,285,794 | 56,193,512 | 84.77 |
| | T3 | 1 | 72,572,070 | 60,028,892 | 82.72 |
| | | 2 | 70,541,562 | 57,902,132 | 82.08 |
| | | 3 | 62,763,430 | 51,703,724 | 82.38 |
| | T4 | 1 | 71,929,368 | 59,468,750 | 82.68 |
| | | 2 | 72,038,762 | 59,670,478 | 82.83 |
| | | 3 | 69,844,624 | 57,754,286 | 82.69 |
| LB | T1 | 1 | 63,596,140 | 53,082,884 | 83.47 |
| | | 2 | 63,533,980 | 52,588,168 | 82.77 |
| | | 3 | 61,307,974 | 51,159,580 | 83.45 |
| | T2 | 1 | 67,996,790 | 56,403,262 | 82.95 |
| | | 2 | 72,179,916 | 60,021,966 | 83.16 |

| | | | | | |
|---|---|---|---|---|---|
| | | 3 | 66,143,426 | 55,361,150 | 83.70 |
| | T3 | 1 | 63,299,082 | 52,946,550 | 83.65 |
| | | 2 | 71,942,830 | 59,202,394 | 82.29 |
| | | 3 | 64,086,828 | 52,905,370 | 82.55 |
| | T4 | 1 | 81,456,084 | 67,291,598 | 82.61 |
| | | 2 | 71,832,870 | 58,985,314 | 82.11 |
| | | 3 | 65,581,214 | 54,881,614 | 83.68 |
| HB | T1 | 1 | 66,529,092 | 54,771,830 | 82.33 |
| | | 2 | 67,687,370 | 56,811,402 | 83.93 |
| | | 3 | 64,823,328 | 53,806,182 | 83.00 |
| | T2 | 1 | 64,300,338 | 54,845,844 | 85.30 |
| | | 2 | 69,442,964 | 57,413,676 | 82.68 |
| | | 3 | 63,179,584 | 52,537,812 | 83.16 |
| | T3 | 1 | 67,012,494 | 54,676,406 | 81.59 |
| | | 2 | 68,906,008 | 56,425,790 | 81.89 |
| | | 3 | 75,585,928 | 59,691,054 | 78.97 |
| | T4 | 1 | 60,510,990 | 50,087,686 | 82.77 |
| | | 2 | 76,408,464 | 63,307,112 | 82.85 |
| | | 3 | 74,420,638 | 61,950,548 | 83.24 |
| VHB | T1 | 1 | 81,001,186 | 67,215,950 | 82.98 |

| | | | | | |
|---|---|---|---|---|---|
| | | 2 | 66,738,486 | 55,112,884 | 82.58 |
| | | 3 | 74,007,774 | 61,992,674 | 83.77 |
| | T2 | 1 | 68,941,894 | 56,603,426 | 82.10 |
| | | 2 | 59,414,832 | 48,526,634 | 81.67 |
| | | 3 | 64,924,224 | 52,769,718 | 81.28 |
| | T3 | 1 | 69,438,956 | 58,202,590 | 83.82 |
| | | 2 | 63,856,034 | 52,275,048 | 81.86 |
| | | 3 | 66,339,142 | 55,246,508 | 83.28 |
| | T4 | 1 | 72,242,966 | 60,019,340 | 83.08 |
| | | 2 | 64,533,060 | 53,713,524 | 83.23 |
| | | 3 | 64,069,950 | 53,694,348 | 83.81 |
| Total | - | - | 3,293,975,098 | 2,729,920,428 | 82.90 |

* VHB: very high ºBrix, HB: high ºBrix; LB: low ºBrix; and VLB: very low ºBrix. T1: 6-month-old plants, T2: 8-month-old plants. T3: 10-month-old plants, T4: 12-month-old plants.
