## Supplementary Table 3 for "Temporal Gene Expression in Apical Culms Shows Early Changes in Cell Wall Biosynthesis Genes in Sugarcane"

**Supplementary Table 3**: Coverage of Sorghum bicolor proteins by the de novo assembled sugarcane transcriptome. Sorghum proteins are binned according to the percentage of coverage by at least one sugarcane transcript.

| Percentage of coverage | Number of proteins matched | Cumulative sum |
|---|---|---|
| 100 | 6.879 | 6.879 |
| 90 | 2.789 | 9.668 |
| 80 | 2.946 | 12.614 |
| 70 | 3.258 | 15.872 |
| 60 | 3.435 | 19.307 |
| 50 | 3.190 | 22.497 |
| 40 | 2.447 | 24.944 |
| 30 | 1.677 | 26.621 |
| 20 | 959 | 27.580 |
| 10 | 201 | 27.781 |
