## Supplementary Data 1 for "Temporal Gene Expression in Apical Culms Shows Early Changes in Cell Wall Biosynthesis Genes in Sugarcane"

**Supplementary Data 1.** Validation of the RNA-Seq expression analysis with RT-qPCR

We validated the RNA-Seq results with RT-qPCR. Gene selection was performed based on the average expression level of genes in the RNA-Seq dataset. We chose differentially expressed genes related to sucrose and cell wall biosynthesis, constitutive genes and genes previously reported in the literature. *Sucrose synthase* and *glyceraldehyde 3-phosphate dehydrogenase* were selected to represent differentially expressed genes. We used the *26S proteasome non-ATPase regulatory subunit 2 homolog gene X2 isoform* (26S) as an internal control.

The primers were designed using Primer3Plus (Untergasser et al., 2007) and Beacon Designer™ Free Edition (http://www.premierbiosoft.com) to avoid secondary structures. The primer design parameters were: fragment size between 90 and 200 bp, primer size ranging from 18 to 23 bp, melting temperature between 50 and 60 °C and GC content from 40 to 60%. We also mapped the primers back to the *de novo* transcriptome and verified if the alignments were unique to isoforms of the same gene.

Synthesis of cDNA was performed with the QuantiTect® Reverse Transcription kit (Qiagen, Chatsworth, CA, USA) using 1000 ng of total RNA. The cDNAs were diluted (1:10) and 2 μl from each sample were used for RT-qPCR. The RT-qPCR assays were conducted with the iTaq Universal SYBR® Green Supermix (Bio-Rad Laboratories Inc., Hercules, CA, USA) following the manufacturer's instructions and using the primer concentration of 0.3 μM. The reactions were performed using the CFX384 Real-Time PCR Detection System with the following cycling conditions: 95 °C for 10 min, followed by 40 cycles of 95 °C at 30 s and 60 °C at 1 min. We used two technical replicates for each biological replicate in the experimental design. The Cq values were estimated with CFX MANAGER 3.1 software (Bio-Rad Laboratories, Inc., USA) and the same contrasts from the RNA-Seq differential expression analysis were tested with the PCR package (Ahmed and Kim, 2018), considering $p < 0.05$. The primers used in this study are described in Table 1, including one pair of primers previously reported in the literature (Hoang et al., 2017).

Table 1: Primers used for RT-qPCR validation. The table contains the corresponding transcript in the *de novo* assembly, its functional annotation and the nucleotide sequence of forward and reverse primers.

| Description | Name | Sequence | Order in pair |
|---|---|---|---|
| *26S proteasome non-ATPase regulatory subunit 2 homolog A isoform X2* | 26S-1 | GTTTCGCCTTTCTCAAGATG | Forward |
| | 26S-2 | ATACGGTTGCTTCCTGTTGC | Reverse |
| *4-coumarate CoA ligase 1* | 4CL1-1 | AGCCGTTCCAGGTCAAGTC | Forward |
| | 4CL1-2 | CTCGGGGTCGTTCAGGTAA | Reverse |
| *Glyceraldehyde 3-phosphate dehydrogenase* (Hoang et al., 2017) | GAPDH-1 | GGTATGTCCTTCCGGGTTCC | Forward |
| | GAPDH-2 | CCACGTAGCCCATGATACCC | Reverse |
| *Sucrose synthase 1* | SuSy1-1 | GTCAAACACATCAATACCAT | Forward |
| | SuSy1-2 | TAACTCTGACCTCTACTGGA | Reverse |
| *Sucrose synthase 4* | SuSy4-1 | GTGAACATCGTGAACTGAGA | Forward |
| | SuSy4-2 | TTATGGTGCTTGGGGTATGC | Reverse |

Four genes were used for validation based on differential expression analysis (Figure 1). The genes *glyceraldehyde 3-phosphate dehydrogenase*, *sucrose synthase 1*, *sucrose synthase 4* and *4-coumarate CoA ligase 1* presented at least 10 comparisons each with matching results between RNA-Seq and RT-qPCR. We performed 58 experiments, of which 46 confirmed the RNA-Seq expression results. In addition, the high correlation between fold changes obtained with both techniques ($r = 0.83$) shows concordance between RNA-Seq and RT-qPCR results (Figure 2).

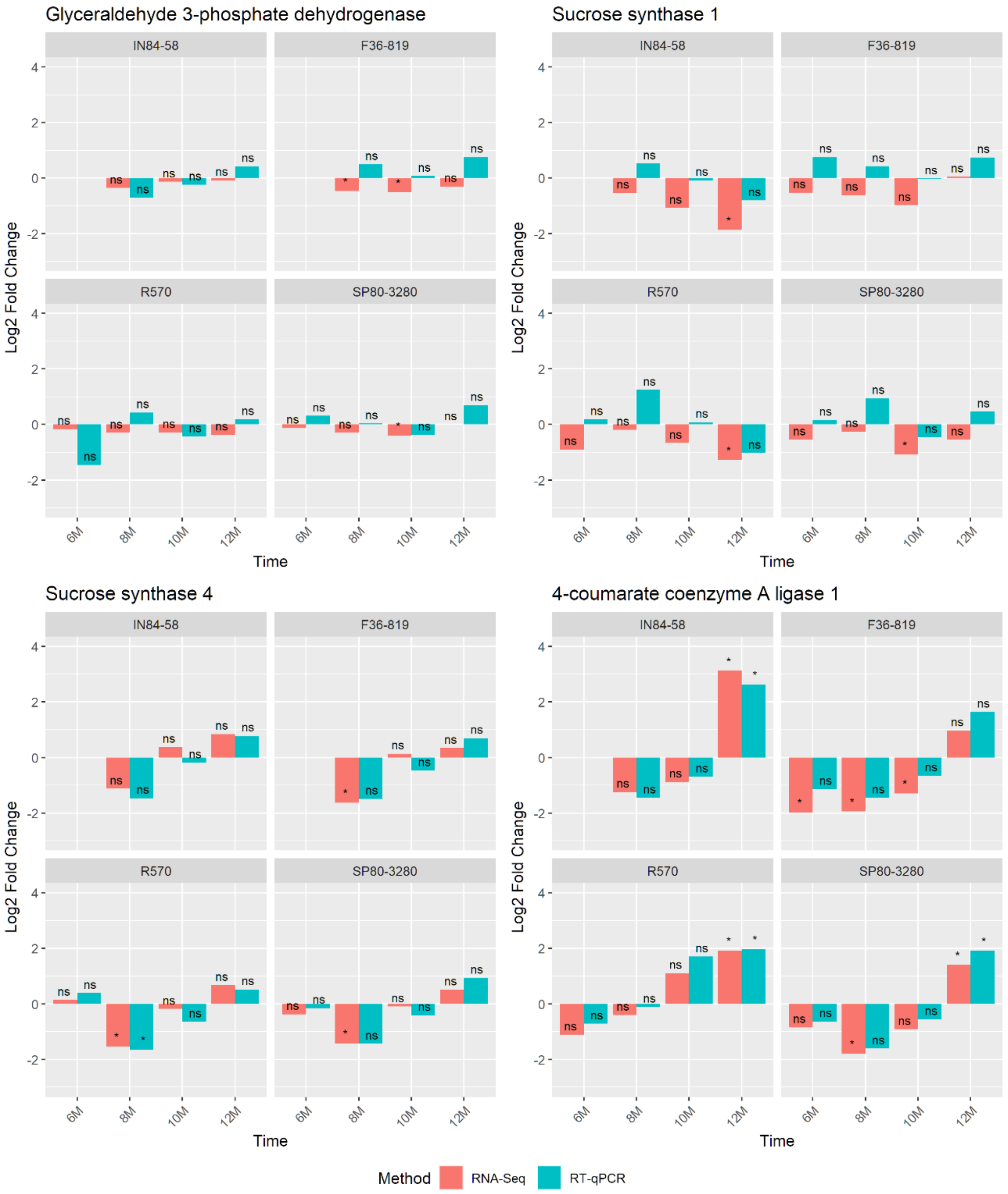


Figure 1: Results of differential expression tests with RNA-Seq and RT-qPCR. The x axis shows the age of plants (in months). The y axis corresponds to the log fold change in comparison to the IN84-58

(very low ºBrix) genotype at six months. Significant tests are indicated with an asterisk, while non-significant ones are marked as ns. Each graph corresponds to a different gene. Red bars indicate the log fold changes from RT-qPCR and blue bars indicate the RNA-Seq log fold changes. The age of plants in the x axis is shown in months.

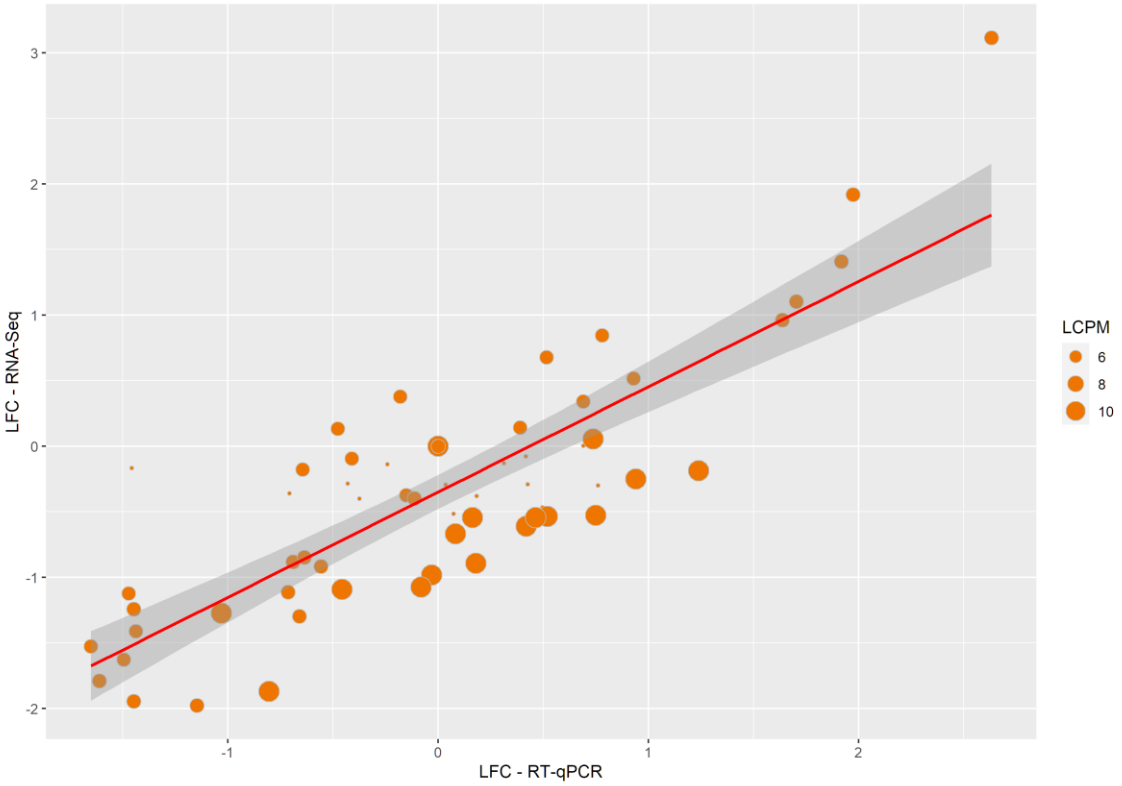


Figure 2: Relationship between log2 fold change (LFC) values obtained with RNA-Seq and RT-qPCR. Each point represents an LFC value for a different gene and genotype, always using IN84-58 in the first time point as a reference. Point sizes indicate the average expression level, measured in log of counts per million (LCPM).
